# Cables2 Is a Novel Smad2-Regulatory Factor Essential for Early Embryonic Development in Mice

**DOI:** 10.1101/744128

**Authors:** Tra Thi Huong Dinh, Hiroyoshi Iseki, Seiya Mizuno, Saori Iijima-Mizuno, Yoko Tanimoto, Yoko Daitoku, Kanako Kato, Yuko Hamada, Ammar Shaker Hamed Hasan, Hayate Suzuki, Kazuya Murata, Masafumi Muratani, Masatsugu Ema, Jun-dal Kim, Junji Ishida, Akiyoshi Fukamizu, Mitsuyasu Kato, Satoru Takahashi, Ken-ichi Yagami, Valerie Wilson, Ruth M. Arkell, Fumihiro Sugiyama

## Abstract

CDK5 and Abl enzyme substrate 2 (Cables2), a member of the Cables family that has a C-terminal cyclin box-like domain, is widely expressed in adult mouse tissues. However, the physiological role of Cables2 *in vivo* is unknown. We show here that *Cables2*-deficiency causes post-gastrulation embryonic lethality in mice. The mutant embryos progress to gastrulation, but then arrest, and fail to grow. Analysis of gene expression patterns reveals that formation of the anterior visceral endoderm and the primitive streak is impaired in *Cables2*-deficient embryos. Tetraploid complementation analyses support the critical requirement of Cables2 in both the epiblast and visceral endoderm for progression of embryogenesis. In addition, we show that Cables2 physically interacts with a key mediator of the canonical Nodal pathway, Smad2, and augments its transcriptional activity. These findings provide novel insights into the essential role of Cables2 in the early embryonic development in mice.

## INTRODUCTION

The Nodal, bone morphogenetic protein (BMP) and Wnt signalling pathways are essential for early mouse development. These pathways coordinately control formation of the proximal– distal (P–D) axis during the egg cylinder stage and the subsequent conversion of this axis into the anterior–posterior (A–P) axis early in gastrulation (reviewed in Arkell and Tam, 2012; Robertson, 2014; Shen, 2007; ten Berge et al., 2008; Wang et al., 2012; Winnier et al., 1995). Initially, ligands of these signalling pathways (including BMP4 and Nodal) and pathway activators (Furin, Pace) are expressed in proximal portions of the embryo whereas antagonists of each pathway (including Cer1, Lefty1, Dkk1) are expressed in the distal-most tissue of the embryo (the distal visceral endoderm: DVE). Subsequently, the distal population of antagonist expressing cells expands and relocates to the anterior of the embryo in a region known as the anterior visceral endoderm (AVE) (Ben-Haim et al., 2006; Shen, 2007). Concurrently, the expression domains of both the Nodal and Wnt3 ligands are confined to the posterior side of the embryo (the primitive streak; PS and overlying visceral endoderm or posterior visceral endoderm PVE). This realignment of the P–D signalling gradient into A–P gradients signals the onset of gastrulation and the formation of the primitive streak, which is the structure that will generate tissues that constitute and elaborate the embryonic A–P axis. The Nodal pathway is crucial for many aspects of A–P axis formation. Mouse embryos that lack any Nodal activity fail to establish molecular pattern in the pre-gastrula VE and lack AVE function. Additionally, these embryos fail to establish posterior identity at the beginning of gastrulation (Brennan et al., 2001; Conlon et al., 1994). Other experiments, in which altered Nodal signalling enables gastrulation to initiate, reveal that Nodal signalling also patterns the primitive streak and its derivatives. High levels of Nodal signalling early in gastrulation ensure the production of the anterior mesendoderm that patterns the embryonic anterior, whereas as gastrulation proceeds the level of Nodal signalling is reduced, ensuring that more-posterior primitive streak derivatives are formed correctly (Vincent et al., 2003). During early mouse development, the Nodal signal is propagated via binding to its receptors activin A receptor, type 1B (Acvr1b, also known as ALK4) and activin A receptor, type 1C (Acvr1c, also known as ALK7). The activated receptors phosphorylate the receptor-regulated Smads (R-Smads; Smad2 and Smad3), which form homomeric complexes and heteromeric complexes with the common Smad (SMAD4). These activated Smad complexes accumulate within the nucleus, where they often complex with tissue specific transcription factors to directly regulate transcription of target genes (Massagué, 2012).

Genetic evidence also implicates the canonical Wnt/β-catenin pathway in multiple aspects of A-P axis formation. Mutations that remove (Huelsken et al., 2000) or increase (Chazaud and Rossant, 2006) Wnt activity prevent correct formation of the DVE. Subsequently, in the 24 hours prior to gastrulation Wnt3 expression is initiated, first being detected in the PVE and then in the posterior epiblast by 6.0 dpc (Rivera-Pérez and Magnuson, 2005). Removal of this activity (Liu et al., 1999) prevents primitive streak formation whereas reduction of this activity (Tortelote et al., 2013; Yoon et al., 2015a) allows the streak to form but not elongate, resulting in impaired production of the tissues that constitute the A-P axis. Generally, when canonical Wnt ligands bind and activate their receptor, the intracellular molecule, β-catenin, is stabilised and translocates to the nucleus where it complexes with the Tcf family of DNA binding proteins to regulate transcription of Wnt target genes (Jamieson et al., 2012; Wang et al., 2012).

The precise level of Nodal and Wnt activity is dependent upon interactions between these and the BMP pathway (Robertson, 2014; Tam and Loebel, 2007). For example, prior to gastrulation, Nodal expression in the epiblast is activated in the overlying visceral endoderm via a Smad2/Foxh1 dependent autoregulatory feedback loop (Norris et al., 2002). At the same time, a slow acting feedback loop is established in which Nodal signals from the epiblast maintain the extraembryonic expression of BMP4 (Ben-Haim et al., 2006; Brennan et al., 2001). BMP4 activates Wnt3 expression in the posterior epiblast and Wnt3 amplifies Nodal expression (Ben-Haim et al., 2006). Although much is known about the signalling events that establish the murine A–P axis, it is clear that many molecules required for this process remain to be discovered.

Cdk5 and Abl enzyme substrate 1 (Cables1, also known as ik3-1) is the founding member of the Cables family, each member of which has a C-terminal cyclin box-like domain. Cables1 has been shown to physically interact with cyclin-dependent kinase 2 (Cdk2), Cdk3, Cdk5, and c-Abl molecules, and to be phosphorylated by Cdk3, Cdk5, and c-Abl (Matsuoka et al., 2000; Yamochi et al., 2001; Zukerberg et al., 2000). Furthermore, in primary cortical neurons, c-abl phosphorylation of Cables1 augments tyrosine phosphorylation of Cdk5 to promote neurite outgrowth (Zukerberg et al., 2000). It has also been demonstrated that Cables1 functions as a bridging factor linking Robo-associated Abl and the N-cadherin-associated β-catenin complex in chick neural retina cells (Rhee et al., 2007). Of note, *Cables1*-deficient mice showed increased cellular proliferation resulting in endometrial hyperplasia, colon cancer, and oocyte development (Kirley et al., 2005; Lee et al., 2007; Zukerberg et al., 2004). Additionally, a dominantly acting, truncated version of Cables1 revealed a requirement in the development of the corpus callosum in mice (Mizuno et al., 2014). During zebrafish development, *Cables1* is required for early neural differentiation and its loss subsequently causes apoptosis of brain tissue and behavioral abnormalities (Groeneweg et al., 2011). Zebrafish have only one Cables gene (Cables1), whereas the mouse and human genomes contain a paralogous gene, Cables2 (also known as ik3-2), which has a C-terminal cyclin-box-like region with a high degree of identity to that of Cables1. Similarly, Cables2 has been shown to physically associate with Cdk3, Cdk5, and c-Abl (Sato et al., 2002). Moreover, forced expression of Cables2 induced apoptotic cell death in both a p53-dependent manner and a p53-independent manner *in vitro* (Matsuoka et al., 2003). The *Cables2* gene is known to be expressed in a variety of adult mouse tissues, including the brain, but the *in vivo* role of this gene has yet to be explored.

To elucidate the role of Cables2 *in vivo*, we generated *Cables2*-deficient mice and found that *Cables2* deficiency caused post-gastrulation embryonic lethality. The mutant embryos progress to gastrulation, but then arrest, and fail to grow. Analysis of gene expression patterns revealed that AVE and PS formation is impaired in *Cables2*-deficient embryos. The expression of both positive and negative components of the Nodal and Wnt signalling pathways are altered at the onset of gastrulation in the mutant embryos. By comparison with many other mouse mutant phenotypes, these defects may be anticipated to give rise either to highly dysmorphic embryos or to embryos with patterning defects. Instead, the *Cables2*-deficient embryos fail to thrive once gastrulation begins but retain the egg cylinder morphology of the early gastrula. This suggests multiple roles for *Cables2* during the immediate post-implantation period. This hypothesis is supported by the ubiquitous expression of Cables2 and by tetraploid complementation analyses which demonstrate that Cables2 functions in visceral endoderm (VE) for AVE and PS formation, whereas epiblast expression of Cables2 regulates embryo growth. We further demonstrated that Cables2 can physically interact with key mediators of the canonical Wnt and Nodal pathways (β-catenin and Smad2 respectively) and augment transcriptional activity of these pathways in cell-based reporter assays. These findings provide novel insights into the essential role of *Cables2* in the early embryonic development in mice.

## RESULTS

### Expression of *Cables2* during early mouse development

*Cables2* is widely expressed at equivalent levels in mouse tissues, including the brain, heart, muscle, thymus, spleen, kidney, liver, stomach, testis, skin, and lung (Sato et al., 2002). We first investigated the expression of *Cables2* in mouse embryonic stem cells (ESCs), blastocysts, and embryos at E7.5 by reverse transcription polymerase chain reaction (RT-PCR). The results indicated that *Cables2* was expressed in all three stages of early development **(Figure 1A)**. To confirm *Cables2* gene expression in mouse embryogenesis, localization of *Cables2* mRNA expression was examined in embryos by WISH **(Figure 1B−F).** The data for the whole embryo and transverse sections showed that *Cables2* was expressed ubiquitously at E6.5 **(Figure 1B and C)**. *Cables2* was detected in both extra- and embryonic parts at E7.5 **(Figure 1D)** and strongly expressed in the allantois and heart-field at E8.5 **(Figure 1E)**. At E9.5, the whole embryo and extraembryonic tissues, including the yolk sac, expressed *Cables2* **(Figure 1F)**. Overall, these data indicate that *Cables2* is expressed ubiquitously during early development, including throughout gastrulation in mouse embryos.

**Figure 1.**
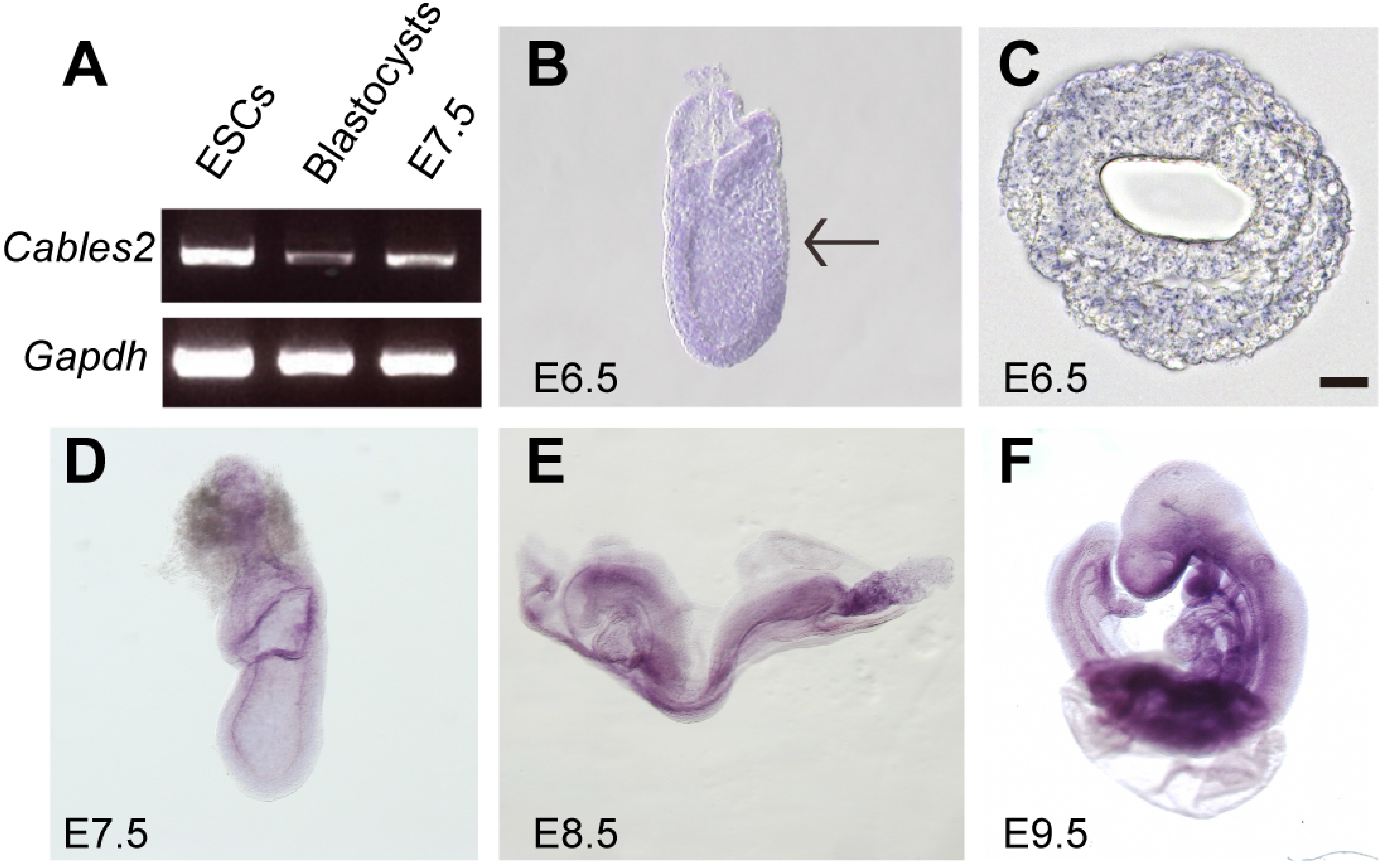
*Cables2* expression during early mouse embryo development. (A) *Cables2* gene expression was examined by RT-PCR with ESC, blastocyst, and E7.5 embryo samples. *Gapdh* was used as an internal positive control. (B–F) Wild-type embryos from E6.5 to E9.5 were examined by *in situ* hybridization with a *Cables2* probe. The whole embryo expressed *Cables2* at E6.5 (B). The black arrow indicates the position of the transverse section shown in (C). Scale bars, 20 μm. The following figure supplement is available for figure 1: **Figure supplement 1.** Genotyping and expression of *Cables2*.

### Early embryonic lethality of *Cables2* deficiency

Next, we generated *Cables2*-deficient mice to investigate the physiological role of Cables2 *in vivo*. *Cables2* heterozygous mice on an inbred C57BL/6N genetic background were produced using conventional aggregation with *Cables2*-targeted ES cell clones. While the heterozygotes were viable and fertile, no homozygous *Cables2*-deficient mice were observed following intercrossing heterozygous mice **(Table 1, Figure 1—figure supplement 1A)**. To identify the critical point in development at which *Cables2* is essential for survival, embryos were collected and genotyped at various time points during embryonic development (**Table 1)**. Homozygous *Cables2* mutant mice were detected in Mendelian ratios at E6.5–E9.5 but no homozygous embryos were observed at or beyond E12.5, indicating that *Cables2*-deficient mice die and are resorbed between E9.5 and 12.5 **(Table 1, Figure 1—figure supplement 1B)**. All of the *Cables2*-deficient embryos collected at E7.5–9.5 were considerably smaller than their wild-type littermates and did not progress beyond the cylindrical morphology of the wild-type early-mid-gastrula **(Table 1, Figure 2A–D)**. Considerably small *Cables2^-/-^* embryos had barely progressed beyond E8.5 **(Figure 2B)**. In section, it was apparent that E7.5 homozygous mutant embryos lacked pattern and instead resembled wild-type embryos in which the primitive streak is just beginning to form (i.e. E6.5 embryos) in both morphology and size **(Figure 2E and F)**. In contrast, all embryos recovered at E6.5 were morphologically indistinguishable from their wildtype littermates **(Table 1)**. Histological analyses confirmed that pre-streak stage (E6.0) *Cables2^-/-^* embryos were structurally normal, exhibiting a normal-sized epiblast, extraembryonic ectoderm, and primitive endoderm **(Figure 2G–H)**. These results suggest that *Cables2* lost-of-function causes growth and patterning arrest early in gastrulation accompanied by post-gastrulation embryonic lethality.

**Table 1.** Survival rate and Mendelian ratio of *Cables2*-mutant embryos.

| Embryonic days (E) | Total number of embryos | Genotypes |  |  |
| --- | --- | --- | --- | --- |
|  |  | +/+ | +/- | -/- |
| E6.5 | 437 | 132 (30.2)<sup>a</sup> | 221 (50.6) | 80 (18.3) |
| E7.5 | 70 | 18 (25.7) | 32 (45.7) | 20<sup>b</sup> (28.6) |
| E8.5 | 21 | 9 (42.9) | 9 (42.9) | 3<sup>b</sup> (14.3) |
| E9.5 | 18 | 7 (38.9) | 7 (38.9) | 4<sup>b</sup> (22.2) |
| E12.5 | 6 | 2 (33.3) | 4 (66.7) | 0 (0) |
| Adult | 90 | 24 (26.7) | 66 (73.3) | 0 (0) |

<sup>a</sup>Number of embryos (percentage), <sup>b</sup>Abnormal phenotype.

**Figure 2.**
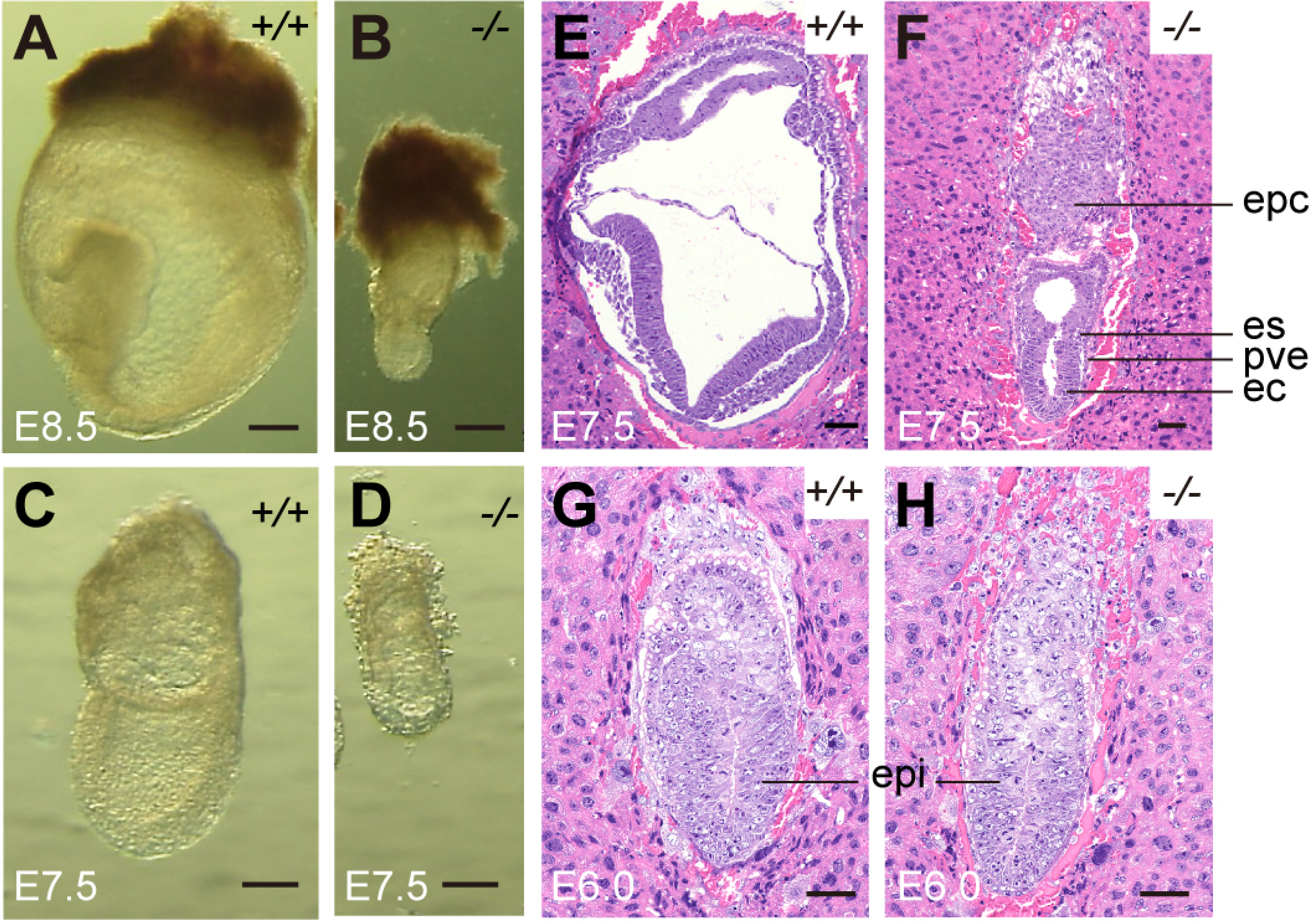
Morphological and histological analyses of *Cables2*-deficient embryos at early stages of development. Embryos were collected and genotyped at E8.5 (A, B) and E7.5 (C, D). Histological analysis was on HE-stained sections. Wild-type and *Cables2* mutant embryos were embedded in paraffin and stained at E7.5 (E, F) and E6.0 (G, H). Epc: ectoplacental cone, ps: primitive streak; pve: posterior visceral endoderm; ec: ectoderm; epi: epiblast. Scale bars, 100 μm (A–D), 50 μm (E–H). The following figure supplement is available for figure 2: **Figure supplement 1**. Proliferating and apoptotic cells in E6.5 *Cables2* mutant embryos.

### Normal cell proliferation and death status in *Cables2*-deficient embryos at E6.5

Cell proliferation and apoptotic cell death are key events during development. To clarify the cell growth status, we performed EdU assay and measured the percentage of EdU-positive cells. There was no significant difference in the percentage of proliferation cells between wild-type and *Cables2^-/-^* embryos at E6.5 **(Figure 2—figure supplement 1A–C)**.

Furthermore, a simultaneous TUNEL assay was performed to determine whether the reduced size of *Cables2*-deficient embryos could be attributed to increased programed cell death. Although apoptotic cells were detected in both the epiblast and embryonic VE, the average percentage of dead cells in *Cables2^-/-^* embryos was not significantly different from that in wild-type embryos **(Figure 2—figure supplement 1D–H)**. These results suggest that cell proliferation and apoptotic cell death are normal in *Cables2*-deficient embryos until E6.5.

### Impaired formation of PS and A–P axis in *Cables2* deficiency

To characterize the phenotype of *Cables2*-deficient embryos, we first analysed the expression of *Brachyury* (*T*). Prior to gastrulation, *T* transcripts are first detected in a ring at the embryonic/extraembryonic junction, whereas once gastrulation is initiated they are found in the PS and nascent mesoderm and subsequently in the axial mesendoderm (Wilkinson et al., 1990). It therefore serves as a marker of the transition from the P-D to A-P axis. At E6.5, *Cables2^-/-^* embryos exhibited normal spatial *T* expression with decreased signal intensity relative to wild-type embryos **(Figure 3A and B)**. By E7.5, *T* transcripts were observed in the PS of the *Cables2^-/-^* embryos, but the transcripts did not extend to the distal point of the embryo and there was no signal in the axial mesendoderm **(Figure 3I and J)**. To confirm the abnormal PS formation in *Cables2^-/-^* embryos, we further investigated the expression of *Fgf8*, a member of fibroblast growth factor family expressed in the PS (Crossley and Martin, 1995), and found that *Fgf8* was decreased in mutant embryos at E6.5 **(Figure 3E and F)**.

**Figure 3.**
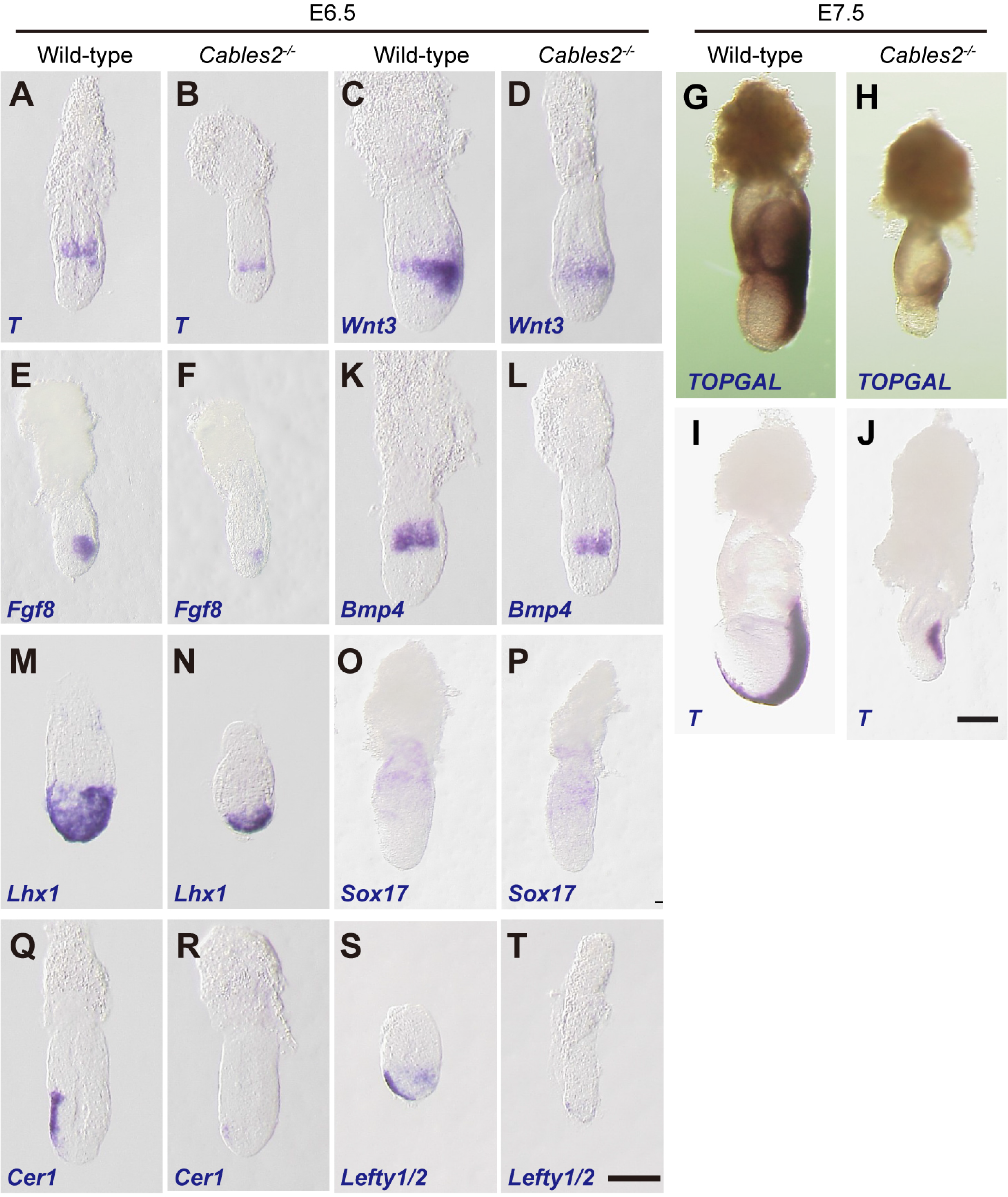
Expression of gastrulation markers in *Cables2*-deficient embryos. (A–F, K–T) All embryos were collected, genotyped, and used for WISH at E6.5. Several key gastrulation markers were examined using both wild-type and *Cables2*-deficient embryos: *T* (*n* = 5), *Wnt3* (*n* = 3), *Fgf8* (*n* = 3), *BMP4* (*n* = 3), *Lhx1* (*n* = 3), *Sox17* (*n* = 3), *Cer1* (*n* = 3) and *Lefty1/2* (*n*= 3). (G, H) β-galactosidase staining demonstrating the restricted activation of Wnt/β-catenin signalling in *Cables2* homozygous embryo carrying the TOPGAL reporter (*n* = 6). (I, J) WISH analysis showing the expression of *T* in wild-type and *Cables2*-deficient embryos at E7.5 (*n* = 5). Scale bars, 100 μm.

PS formation and progression is dependent upon canonical Wnt signalling driven by the expression of *Wnt3* in the proximal-posterior epiblast and PVE (Mohamed et al., 2004; Yoon et al., 2015a) and *T* is a direct target of this Wnt activity (Arnold et al., 2000). WISH showed that expression of *Wnt3* was also impaired in the proximal-posterior part of epiblast and the PVE of E6.5 *Cables2^-/-^* mutants although the expression remained in the proximal epiblast adjacent to the extraembryonic ectoderm (ExE) **(Figure 3C and D)**. To assess the functional significance of altered *Wnt3* expression, *Cables2*-deficient animals were crossed with the TOPGAL transgenic mice, which express the β-galactosidase under the control of three copies of the Wnt-specific LEF/TCF binding sites (Moriyama et al., 2007). In wild-type E7.5 embryos carrying TOPGAL, the β-galactosidase was detected in the fully elongated primitive streak and in the adjacent posterior tissues as expected **(Figure 3G)**. In contrast, E7.5 *Cables2^-/-^* embryos carrying TOPGAL showed the weak expression of β-galactosidase only in proximal-posterior PS **(Figure 3H)**. Meanwhile, *Bmp4* is known to be expressed in the ExE of the post-implantation mouse embryo where it promotes *Wnt3* expression in the proximal epiblast for PS formation. WISH analyses showed that *Bmp4* was similarly expressed in the ExE of *Cables2^-/-^* embryos compared with wild-type embryos at E6.5 **(Figure 3K and L)**, suggesting that the ExE is normally developed in mutant embryos at least until E6.5. These findings are consistent with *Cables2* promoting the formation of *Wnt3*-expressing PVE to induce and maintain PS formation.

Given that apparent impairment of proximal/posterior development in the *Cables2^-/-^* embryos, we next examined markers of the distal/anterior components of the axis. *Lhx1*, which is normally expressed in the AVE and nascent mesoderm of wild-type embryos, was accumulated in the distal part of E6.5 *Cables2*-deficient embryos **(Figure 3M and N)**, whereas the normal formation of extraembryonic VE in mutant embryo was confirmed by the expression of *Sox17* **(Figure 3O and P)**. Our data also showed that *Cerberus 1* (*Cer1*) and *Lefty1*, antagonists of Nodal signalling, were expressed at lower levels in *Cables2^-/-^* embryos compared to the wild-type at E6.5 **(Figure 3Q–T)**. Furthermore, WISH analyses demonstrated absent or decreased expression of *Lefty2* in the posterior epiblast of *Cables2*-deficient embryos at E6.5 **(Figure 3S and T)**. Taken together, the results of WISH analyses suggest that *Cables2* depletion impairs the correct AVE formation at the gastrulation stage.

### Activation and interaction of Cables2 with β-catenin

Accumulating evidences have suggested that Wnt/β-catenin signalling is implicated in the formation of AVE (Engert et al., 2013; Huelsken et al., 2000; Lickert et al., 2002) and it is known that the Cables2 paralog (Cables1) can bind to β-catenin (Rhee et al., 2007). We therefore examined whether Cables2 facilitates β-catenin activity at Wnt target sites and physically interacts with β-catenin. The results indicated that Cables2 activated β-catenin/TCF-mediated transcription *in vitro* with an almost two-fold increase in relative TOP/FOP luciferase activity **(Figure 4A)**. Moreover, co-IP using N-terminal FLAG-tagged Cables2 (FLAG-Cables2)-transfected 293FT cell lysates with or without exogenous β-catenin indicated that β-catenin was present in the precipitated complexes with Cables2 **(Figure 4B and Figure 4—figure supplement 1)**. These data suggest that Cables2 can physically associated with β-catenin and increases its transcriptional activity at Wnt-responsive genes.

**Figure 4.**
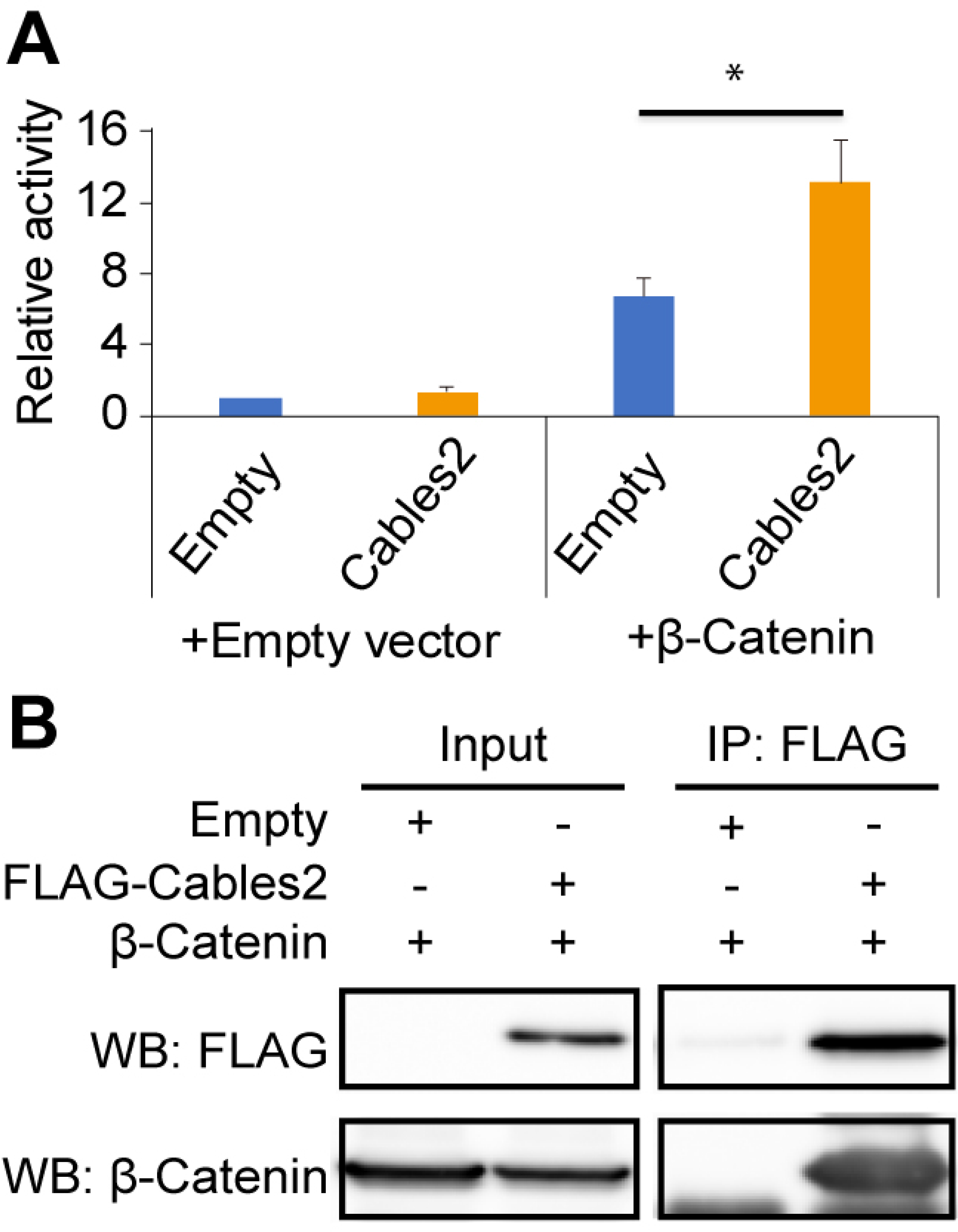
Enhancement of β-catenin activity by Cables2. (A) Relative luciferase activities in 293T cells transfected with an empty control or Cables2 expression vectors together with an empty control or β-catenin expression vectors. Relative luciferase activity is expressed as the ratio of TOP/FOPflash reporter activity relative to the activity in cells transfected with an empty vector alone. Columns: Averages of at least three independent experiments performed in triplicate. Error bars, Standard deviation (SD). Statistical significance was determined using Student’s *t* test (*, *P* < 0.05). (B) Co-IP was performed with FLAG-Cables2 and β-catenin expression vectors. The results obtained using anti-FLAG and anti-β-catenin antibodies showed the appearance of β-catenin in the precipitated complexes with Cables2. The following figure supplement is available for figure 4: **Figure supplement 1.** Interaction of Cables2 with endogenous β-catenin.

### Activation and interaction of Cables2 with Smad2

Beside β-catenin, proper activation of the Nodal/Smad2 signalling in VE is required for the AVE formation (Takaoka and Hamada, 2012). Thus, we conducted *in vivo* and *in vitro* experiments to examine whether Cables2 impacts Nodal/Smad2 signalling. WISH analysis revealed the normal expression of *Nodal* in *Cables2^-/-^* embryos at E6.0 **(Figure 5A and B)**. Subsequently *Nodal* expression normally localizes to the nascent PS and the posterior epiblast at E6.5, however, in E6.5 *Cables2-*deficient embryos *Nodal* expression remains throughout the epiblast **(Figure 5C and D)**. To determine whether this mislocalisation of *Nodal* expression altered Anterior axis formation, we examined expression of *Foxa2*, a downstream target of Nodal/Smad2 signalling (Brennan et al., 2001; Liu et al., 2004), which is an early marker of Anterior axis formation and necessary for mesendodermal specification (Dufort et al., 1998). As the result, *Foxa2* expression is downregulated in E7.5 *Cables2^-/-^* embryos **(Figure 5E and F**). In addition, we examined the expression of neuroectoderm markers *Sox2* and *Otx2*. WISH revealed that the localized expression of *Sox2* to the anterior embryonic part was decreased in *Cables2^-/-^* embryos **(Figure 5G and H)**, whereas *Otx2* was ectopically expressed in the posterior region of *Cables2^-/-^* embryos at E7.5 **(Figure 5I and J**). These results, in conjunction with the AVE analyses, support our notion that loss of *Cables2* also impairs distal/anterior axis formation and suggest that Cables2 is involved in reinforcement of Nodal/Smad2 signalling in the AVE.

**Figure 5.**
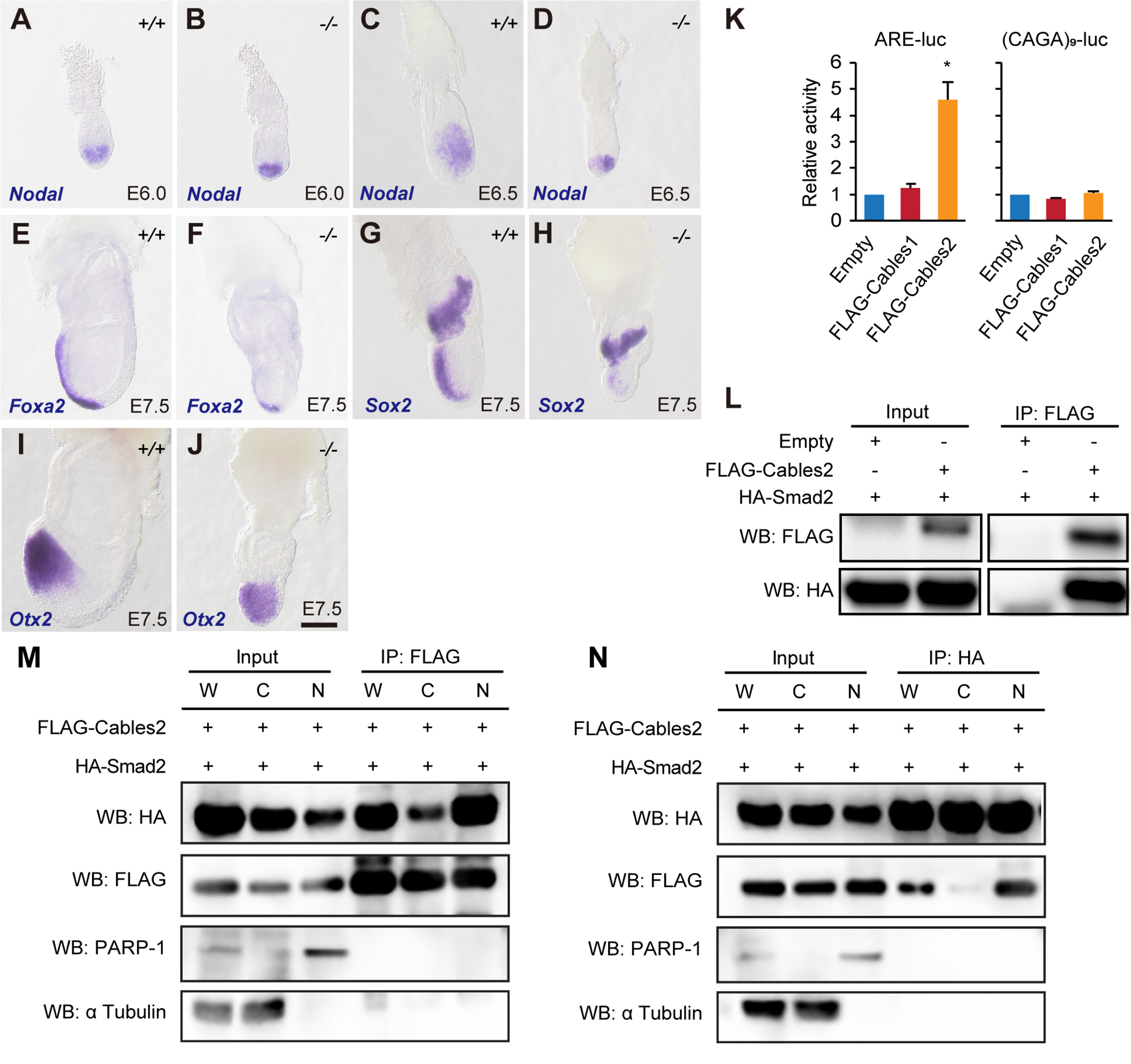
Enhancement of Smad2 activity by Cables2. (A–J) WISH analyses showing expression of *Nodal* (E6.0: *n* = 3; E6.5: *n* = 4), *Foxa2* (*n* = 3), and neuroectoderm markers *Sox2* (*n* = 4) and *Otx2* (*n* = 5). Scale bars, 100 μm. (K) Relative luciferase activities of ARE-luc or (CAGA)_9_-luc reporter vectors in 293FT cells co-transfected with an empty control, FLAG-Cables1, or FLAG-Cables2 expression vectors. Columns: Averages of at least three independent experiments performed in triplicate. Error bars: Standard deviation. Statistical significance was determined using Student’s *t* test (*, *P* < 0.05). (L) Co-IP showing the physical interaction of FLAG-Cables2 with HA-Smad2. (M, N) Co-IP/western blot analysis performed with whole cell lysates or fractionated lysates from 293FT cells transfected with HA-Smad2 and either FLAG-Cables2 or empty control vectors. PARP-1 and αTubulin serve as a nuclear and cytoplasmic marker, respectively. W: whole cell lysate, C: cytoplasmic fraction, N: nuclear fraction. The following figure supplement is available for figure 5: **Figure supplement 1.** Interaction of Cables2 with endogenous Smad2.

We next investigated whether Cables2 enhanced the transcriptional activity of Smad2 by luciferase reporter assay with the ARE-luc vector, which expresses firefly luciferase in a Smad2- and FAST1-dependent manner. We co-transfected 293FT cells with the ARE-luc, FAST1 expression vectors and a constitutive-active mutant form of TGF-β receptor together with an empty vector or FLAG-Cables2 expression vector. The data indicated that forced expression of Cables2 facilitated Smad2 activity in 293FT cells **(Figure 5K)**. Intriguingly, Cables2 had no effect on luciferase activity of the Smad3/4-specific reporter vector (CAGA)_9_-luc (Dennler et al., 1998) **(Figure 5K)**, suggesting that Cables2 does not function as an activator for Smad3. Luciferase assays also revealed that Cables1 had no effect on the transcriptional activities of both Smad2 and Smad3 **(Figure 5K)**. These findings suggest that the promoting activity on Smad2 function is a unique property of Cables2 rather than a conserved function of the Cables family. We further conducted co-IP experiments of Cables2 with exogenous and endogenous Smad2 and found that Cables2 physically interacted with Smad2 in 293FT cells **(Figure 5L–N and Figure 5—figure supplement 1)**. Moreover, co-IP experiments with fractionated extracts demonstrated that FLAG-Cables2 and HA-Smad2 were precipitated from both cytoplasmic and nuclear extracts (**Figure 5L–N**), suggesting that Cables2 forms a complex with Smad2 in both cytoplasm and nucleus. These results suggest that Cables 2 can act as a positive regulatory factor of Smad2.

### Facilitation of *Nanog* expression and its promoter activity by *Cables2*

To further investigate a functional relationship between Cables2 and Smad2 in epiblast, we established *Cables2*-deficient ESCs from homozygous embryos at E3.5 and induced their differentiation into epiblast-like cells (EpiLCs). The morphology, proliferation, and expression of pluripotency genes in *Cables2*-deficient ESCs were similar to those in wild-type ESCs **(Figure 6—figure supplement 1A)**. Both types of ESCs exhibited similar morphological changes after EpiLC induction with activin and bFGF **(Figure 6—figure supplement 1B)**. Nanog plays a crucial role in early mouse embryonic development. The expression of *Nanog* has been observed in the cells of the inner cell mass (ICM) of the E3.5 blastocyst and the epiblast in the egg cylinder at the PS stage (Chambers et al., 2003; Hatano et al., 2005). The cytokine dependency of *Nanog* expression is known to switch from LIF/Stat in the ICM to Nodal/Smad2 in the epiblast. It is noteworthy that *Nanog* expression is highly dependent on Smad2 but not on Smad3 (Sakaki-Yumoto et al., 2013; Sun et al., 2014). Since our luciferase assay revealed that Cables2 functioned as an activator specific for Smad2 **(Figure 5K)**, we focused on and analysed the expression level and promoter activity of *Nanog* in EpiLCs lacking *Cables2*. Interestingly, quantitative RT-PCR showed that *Nanog* mRNA level was decreased by approximately 40% in *Cables2*-deficient EpiLCs **(Figure 6A)**. Moreover, luciferase assay demonstrated that *Cables2* deficit reduced *Nanog* promoter activity in EpiLCs **(Figure 6B)**. Intriguingly, Cables2 physically interacted with Oct4 that is a transcription factor activating *Nanog* and PS gene promoters synergistically with Smad2 **(Figure 6—figure supplement 2)**. Together with the previous findings **(Figure 5K and L)** (Funa et al., 2015; Sakaki-Yumoto et al., 2013; Sun et al., 2014), the results suggest that Cables2 positively regulates *Nanog* expression via the interaction with Smad2 and Oct4 in EpiLCs.

**Figure 6.**
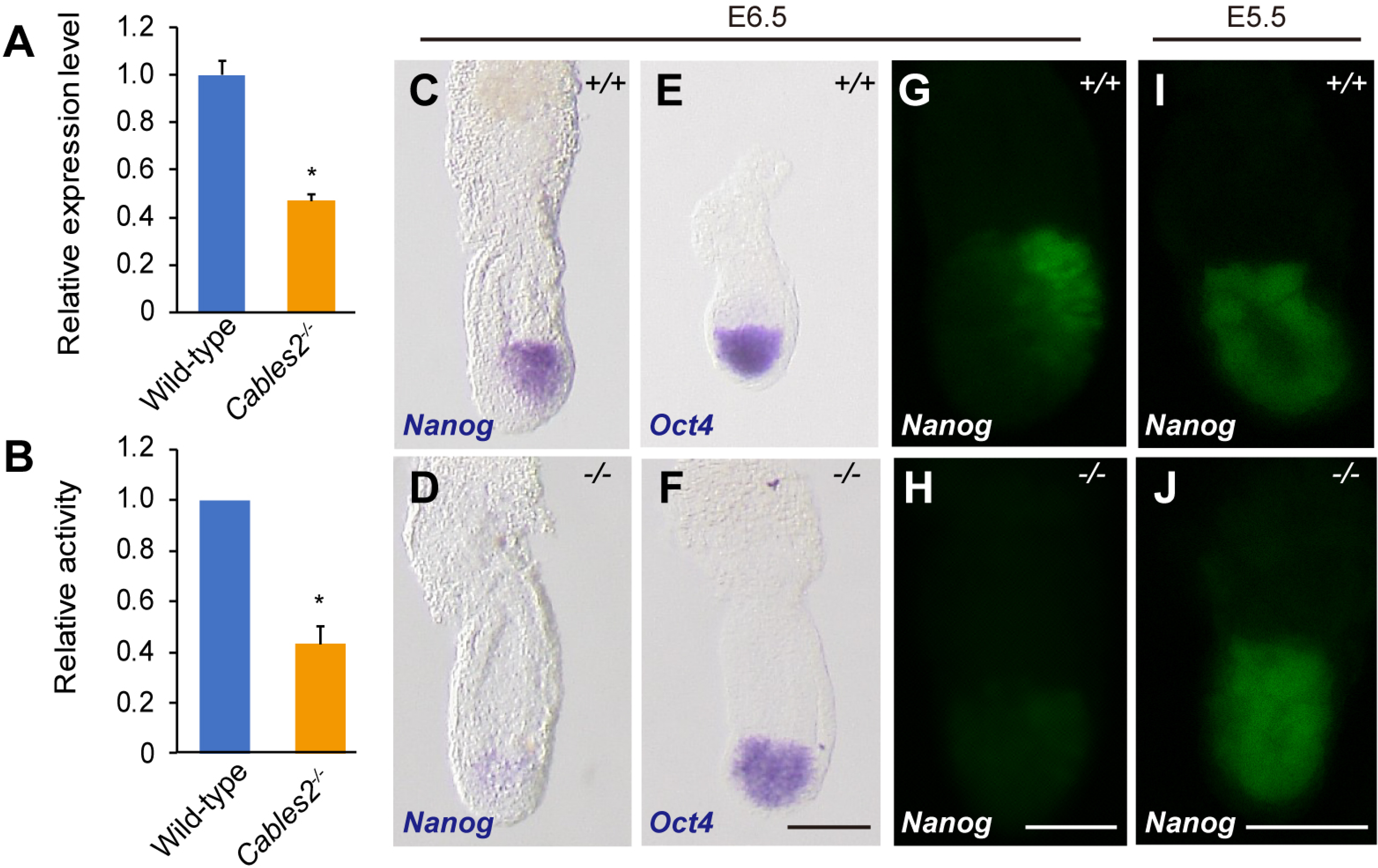
Downregulated *Nanog* promoter activity and expression. EpiLCs were induced from homozygous *Cables2* mutant ESCs. RT-qPCR (A) and *Nanog*-luc assay (B) consistently showed decreased expression of *Nanog* in *Cables2*-deficient cells compared with wild-type EpiLCs. (C, D) *Nanog* expression in *Cables2* mutants was weaker and located evenly in the epiblast compare with wild-type on WISH analysis (*n* = 4). (E, F) *Oct4* was expressed normally in epiblast of mutant embryos (*n* = 4). (G, H) All embryos in the same litter were collected at E6.5, and those with GFP-positive cells were compared. *Cables2^-/-^* embryos showed faint and ectopic expression of the *Nanog*-GFP signal. (I, J) There was no difference between wild-type and mutant embryos at E5.5. Columns: Averages of at least three independent experiments performed in triplicate. Y error bars: Standard deviation (SD). Statistical significance was determined using Student’s *t* test (*, *P* < 0.05). Scale bars, 100 μm (C–H); 50 μm (I, J). The following figure supplements are available for figure 6: **Figure supplement 1.** EpiLC induction from *Cables2^-/-^* ESCs and expression of pluripotency genes. **Figure supplement 2.** Interaction of Cables2 with exogenous HA-Oct4. **Figure supplement 3.** Gene enrichment analyses of up-regulated genes by the loss of Cables2. **Figure supplement 4.** Gene enrichment analyses of down-regulated genes by the loss of Cables2.

To gain insight into changes in global gene expression in *Cables2*-deficient EpiLCs, we performed RNA-seq and gene enrichment analyses for Gene ontology (GO) terms and Kyoto Encyclopedia of Genes and Genomes (KEGG) pathways using the DAVID Bioinformatics Resources (Huang et al., 2009a; Huang et al., 2009b). Notably, the analyses showed GO term enrichments related to “nervous system development” and “negative regulation of cell proliferation” and KEGG pathway enrichments related to several signaling pathway including “p53 signalling pathway” among 122 out of 125 upregulated genes in *Cables2*-deficient EpiLCs **(Figure 6—figure supplement 3A and B)**. While, 52 out of 59 downregulated genes in *Cables2*-deficient EpiLCs represented GO terms related to “proximal/distal pattern formation”, “positive regulation of cell proliferation”, and “male gonad development” and KEGG pathway related to “signalling pathways regulating pluripotency of stem cells” **(Figure 6—figure supplement 4A and B)**.

Previous studies have addressed *Nanog* expression in mouse embryos at early embryonic stages by immunostaining and WISH. The data indicated that *Nanog* is expressed in the whole region of E5.5 epiblasts, but only in the posterior region of E6.5 and E7.5 epiblast (Hart et al., 2004; Hatano et al., 2005). Consistent with these previous reports, our WISH data showed that *Nanog* was expressed in the posterior region of wild-type embryos at E6.5 **(Figure 6C)**. In contrast, *Cables2*-deficient embryos showed only a low level of *Nanog* gene expression over the whole epiblast region at E6.5 **(Figure 6D)**. On the other hand, the pluripotency marker, *Oct4*, was expressed normally in *Cables2^-/-^* mutants at E6.5 (**Figure 6E and F**). To further confirm the expression pattern of *Nanog* in the E6.5 embryo, we utilized *Nanog*-GFP transgenic mice (Okita et al., 2007). Mice carrying the *Nanog*-GFP reporter were crossed with *Cables2*-deficient heterozygotes to obtain *Cables2^-/-^* reporter embryos. At E6.5, *Cables2^-/-^* embryos showed no elevation of *Nanog*-GFP expression at the PS **(Figure 6G and H)**. Nevertheless, there was no difference in GFP expression between embryos of the same litter at E5.5 **(Figure 6I and J)**, suggesting no detectable phenotype of *Nanog* before PS formation. Overall, loss of *Cables2* leads to downregulated *Nanog* expression at the start of gastrulation.

### Requirement of *Cables2* in both epiblast and VE for the proper gastrulation in mice

To determine whether Cables2 is required in the VE, the epiblast, or both, chimera analysis was performed using tetraploid wild-type embryos and *Cables2^-/-^* ESCs. In tetraploid complementation chimera has the advantage that the host tetraploid embryos can only contribute to primitive endoderm derivatives and trophoblast compartment of the placenta, whereas epiblast components are completely derived from ESCs (Tanaka et al., 2009). Tetraploid wild-type morula was aggregated with *Cables2^-/-^* ESCs to produce chimera in which the *Cables2* was exclusively deleted in epiblast but not in the VE (*Cables2* VE rescue chimera) **(Figure 7A)**. *YFP* reporter gene was inserted into *ROSA26* locus of *Cables2^-/-^* ESC to construct *Cables2^-/-^*; *ROSA26^YFP/+^* ESC which gives the advantage for embryo visualisation and imaging. We collected the *Cables2* chimeric embryos at the indicated embryonic days and analysed the phenotype **(Table 2)**. Like *Cables2^-/-^* embryos, epiblast of *Cables2* VE rescue embryos were smaller in size than that of control wild-type chimera littlemates at E7.5 and E8.5 **(Figure 7B–E)**. However, *T* and *Foxa2* were properly expressed in the posterior epiblast and the anterior midline mesendoderm of E7.5 *Cables2* VE rescue embryos, respectively **(Figure 7F and G)**, suggesting that the embryos with *Cables*2^-/-^ epiblast and wild-type VE can form the PS and A–P axis. Importantly, some *Cables2* VE rescue chimeras developed up to E8.5 exhibited the clear structure of head-fold, node and tail bud although embryo size was still extremely smaller compared with wild-type chimeras **(Figure 7H)**. These results suggest that *Cables2* in the VE plays an essential role in the formation of the PS and AVE while Cables2 in the epiblast is essential for epiblast growth.

**Table 2.** Phenotypes in Cables2 VE rescue and Cables2 VE & Epi rescue chimeras.

|  | Tetraploid embryo + *Cables2*<sup>-/-</sup>;<br>*ROSA26*<sup>YFP/+</sup> ESC<br>(VE rescue chimera) |  |  | Wild-type chimera |  |  |
| --- | --- | --- | --- | --- | --- | --- |
| Embryonic<br>days (E) | Total number<br>of embryos | Phenotype |  | Total<br>number of<br>embryos | Phenotype |  |
|  |  | Normal | Abnormal |  | Normal | Abnormal |
| E6.5 | 2 | 1 | 1 | 4 | 3 | 1 |
| E7.5 | 15 | 4 | 11 | 14 | 11 | 3 |
| E8.5 | 13 | 0 | 13<sup>a</sup> | 7 | 6 | 1 |
|  | Tetraploid embryo + *Cables2*<sup>-/-</sup>;<br>*ROSA26*<sup>YFP/+</sup>; *CAG-tdTomato-2A-*<br>*Cables2-3xFLAG* ESC<br>(VE & Epi rescue chimera) |  |  | Wild-type chimera |  |  |
| Embryonic<br>days (E) | Total number<br>of embryos | Phenotype |  | Total<br>number of<br>embryos | Phenotype |  |
|  |  | Normal | Abnormal |  | Normal | Abnormal |
| E7.5 | 5 | 4 | 1 | 2 | 1 | 1 |
| E8.5 | 4 | 3 | 1 | 5 | 4 | 1 |
| E9.5 | 2 | 2 | 0 | 3 | 2 | 1 |

<sup>a</sup>All embryos had A-P axis specification.

**Figure 7.**
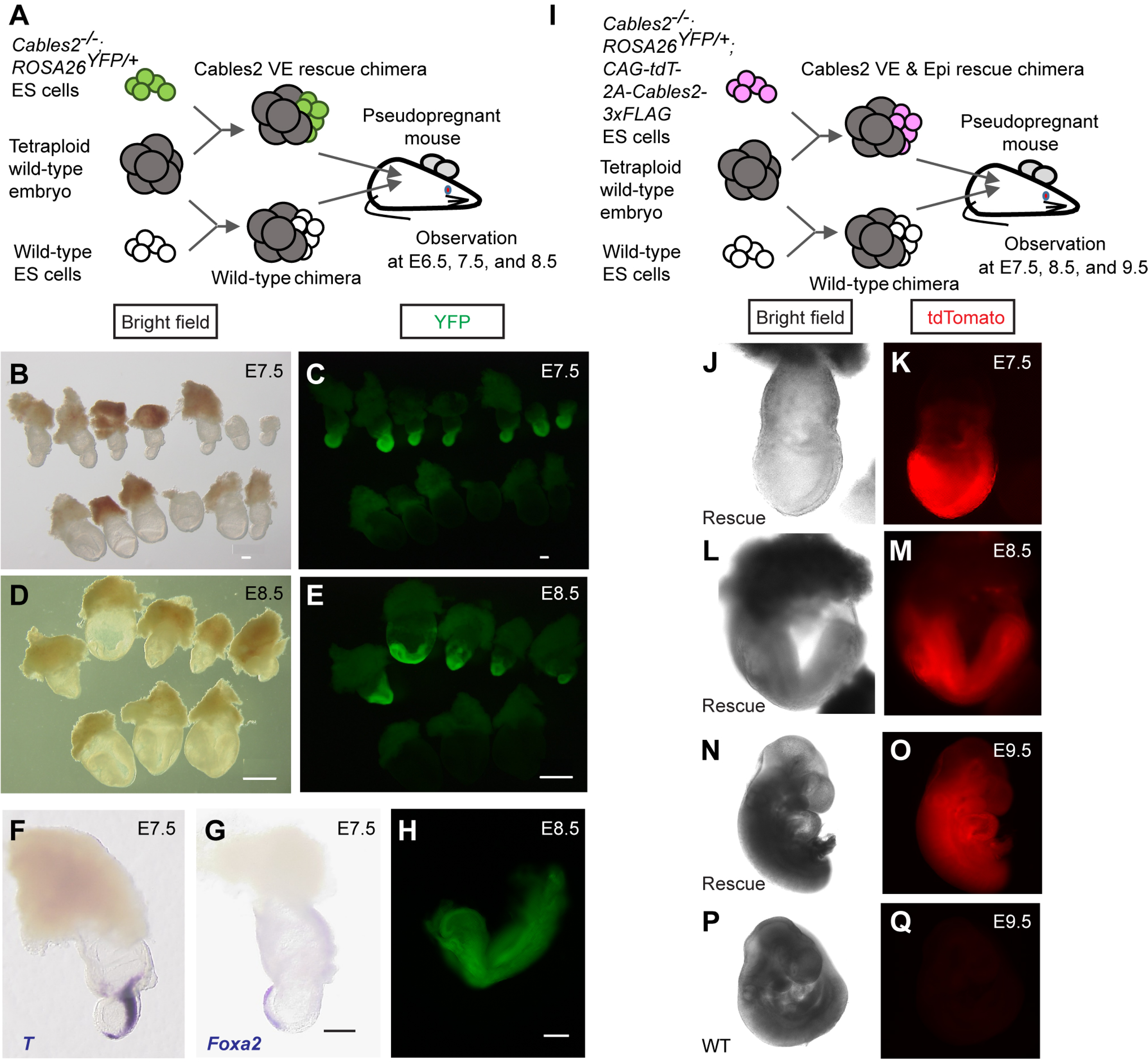
A–P axis formation in *Cables2* VE rescue and normal phenotype of *Cables2* VE & Epi rescue chimeras. (A) Schematic diagram of tetraploid complementation experiment for *Cables2* VE rescue chimera. (B–E) Bright field (B, D) and YFP fluorescent (C, E) images of wild-type and *Cables2* VE rescue chimeric embryos at E7.5 or E8.5. *Cables2* VE rescue chimeras at E8.5 showed distinguishable phenotypes in compared with wild-type chimeras. (F, G) WISH analyses showing the expression of *T* (F) and *Foxa2* (G) in *Cables2* VE rescue chimeric embryos at E7.5. (H) Representative image of *Cables2* VE rescue chimeric embryo at E8.5 showing the head-fold and allantois bud formation. (I) Schematic diagram of tetraploid complementation experiment for *Cables2* VE and epiblast (*Cables2* VE and Epi) rescue chimeras. (J–Q) Bright field (J, L, N, P) and tdTomato fluorescent (K, M, O, Q) images of wild-type and *Cables2* VE and Epi rescue chimeric embryos at E7.5, E8.5, or E9.5. The *Cables2* VE and Epi rescue chimeric embryos developed normally until E9.5. Scale bars, 100 μm (B, C, G, H); 500 μm (D, E).

To confirm the essential function of *Cables2* in epiblast, tetraploid wild-type morula was aggregated with *Cables2^-/-^*; *ROSA26^YFP/+^*; *CAG-tdTomato-2A-Cables2* ESCs to produce *Cables2* VE and epiblast rescue chimera (*Cables2* VE & Epi rescue) (**Figure 7I**). *CAG-tdTomato* fused *2A-Cables2* was randomly integrated into the genome of *Cables2^-/-^ ROSA26^YFP/+^* ESC to ubiquitously overexpress Cables2 in epiblast and its derivatives. Red fluorescence was detected in *Cables2^-/-^*; *ROSA26^YFP/+^*; *CAG-tdTomato-2A-Cables2* ESCs, suggesting that tdTomato-2A-Cables2 was correctly translated in the cells (**Figure 7J–Q**). As the result, *Cables2* VE & Epi rescue chimeras were indistinguishable from wild-type chimera littermates at all time points examined (**Figure 7J–Q**). Of note, the lethal phenotype of *Cables2^-/-^* embryos during gastrulation was rescued by the exogenously expressed Cables2. Altogether, the findings from chimera experiments indicate that Cables2 in both epiblast and VE is required for early embryonic development in mice.

## DISCUSSION

In this study, we provided the first evidence regarding the physiological roles of Cables2 in mice. We demonstrated that *Cables2* is expressed ubiquitously during early embryonic development and that disruption of the *Cables2* gene caused defective A–P axis formation, growth retardation and post-gastrulation embryonic lethality. Many other mouse mutants with impaired A–P axis formation either become highly dysmorphic, or complete gastrulation without growth retardation and instead exhibit patterning defects. The divergent phenotype that results from *Cables2* deficiency therefore suggests it may play multiple roles during early mouse development. Tetraploid complementation experiments demonstrated that the A–P axis and growth phenotypes can indeed be separated. The defective A–P axis formation can be attributed to a requirement for *Cables2* expression in the VE of the mouse embryo, whereas the growth retardation is caused by loss of *Cables2* function in the epiblast. Similarly, at the molecular level Cables2 may play multiple roles. We show here that Cables2 can interact with pivotal intracellular components of both the Nodal and Wnt pathways.

*Cables2*-deficient embryos show a defect in the establishment of the AVE as judged by the downregulation of *Cer1* and *Lefty1* in this structure at E6.5 (**Figure 3Q–T**). The expression of these genes, as well as other aspects of AVE formation, are Nodal/Smad2 dependent (Brennan et al., 2001; Conlon et al., 1994; Kumar et al., 2015; Nomura and Li, 1998; Waldrip et al., 1998; Weinstein et al., 1998). Notably, chimeric embryo experiments have revealed that the Nodal pathway is required in the VE (but not epiblast) to ensure AVE production (Brennan et al., 2001). Moreover, similar experiments have shown that, in the VE, Smad2 alone transduces the Nodal signal required for AVE formation (Heyer et al., 1999; Waldrip et al., 1998) consistent with the fact that Smad3 is not expressed in this tissue (Tremblay et al., 2000). In contrast, Smad3 can compensate for the role of Smad2 in Nodal signalling in the epiblast (Brennan et al., 2001; Dunn et al., 2005; Tremblay et al., 2000). The chimeric experiments conducted here show that, like Nodal and Smad2, Cables2 is also required in the VE to ensure correct AVE formation (**Figure 7A–H**). Furthermore, we demonstrated that Cables2 can physically associated with Smad2 and enhanced the Smad2/FAST1-mediated transcription when overexpressed in cells (**Figure 5K and L**). During murine development, Nodal expression in the VE (but not epiblast) utilises an intronic enhancer that is Smad2/FAST1 dependent (Norris et al., 2002). Overall, these data are consistent with Cables2 interacting with the Nodal/Smad2 pathway in the VE to correctly establish the AVE.

In embryos deficient for *Cables2*, Nodal expression does not become upregulated at the prospective posterior of the embryo (**Figure 5C and D**). At this stage of normal development, Nodal signals from the epiblast are required to promote the expression of posterior genes such as *Wnt3* and *T*, both of which are downregulated in *Cables2*-deficient embryos, as is TCF-mediated transcription measured by the TOPGAL reporter mouse (**Figure 3A–D, G, H**). This posterior Wnt/β-catenin activity is required for primitive streak formation and elongation. The truncated primitive streak and derivatives seen in the *Cables2* mutant embryos (**Figure 3I, J and 5E, F**) are therefore consistent with the decreased posterior Wnt activity in these embryos. Both of these aspects of the phenotype are rescued when *Cables2* is expressed in the VE alone, implying that in these embryos Wnt activity is elevated to a level consistent with relatively normal streak formation and elongation. It is possible that the VE source of *Cables2* elevates the posterior Wnt/β-catenin activity directly (via Cables2/β-catenin interaction), or indirectly via the Nodal/Smad2 pathway and associated upregulation of *Wnt3* expression.

Genetic experiments that ablate *Wnt3* activity in either the VE or epiblast alone (and therefore reduce overall posterior Wnt signal) have shown that this activity regulates the timing of primitive streak formation. Embryos in which VE *Wnt3* function is ablated have delayed primitive streak formation but by E9.5 are indistinguishable from wild-type littermates (Yoon et al., 2015a). When *Wnt3* function is removed from the epiblast, primitive streak formation is delayed and by mid-gastrulation the embryos are highly dysmorphic with the primitive streak bulging towards the amniotic cavity (Tortelote et al., 2013). In contrast, *Cables2*-deficient embryos in which *Wnt3* expression and activity is reduced exhibit primitive streak abnormalities but neither recover nor become highly dysmorphic. Instead, they retain an egg-cylinder morphology and fail to grow. This suggests that the growth failure of the *Cables2* is not caused by improper A–P axis specification and instead represents a separate function of *Cables2*. This hypothesis is supported by the chimeric experiments that demonstrate that *Cables2* expression in the VE is able to rescue the A–P axis, but not the growth defects.

The underlying molecular cause of the growth defect in the *Cables2*-deficient embryos remains unclear. We confirmed the requirement of *Cables2* in epiblast for proper growth by tetraploid complementation analysis (**Figure 7I–Q**). Nodal signalling has shown to be required for sustaining epiblast proliferation during egg cylinder stage in mice (Stuckey et al., 2011; Yang et al., 1998). However, it is unlikely that the growth retardation of *Cables2*-deficient embryos is associated with the Nodal/Smad2 signalling because epiblast size is normal in *Smad2*-deficient mice due to functional compensation by Smad3 (Brennan et al., 2001; Dunn et al., 2005; Tremblay et al., 2000). An alternative possibility is that Cables2 may function with other factors, e.g. BMP signalling-related proteins that is known to be essential for epiblast proliferation (Mishina et al., 1995). However, proliferative defects were apparent in the E6.5 embryos that lack the Bmpr1 receptor (Mishina et al., 1995), unlike in the *Cables2*-deficient embryos where no alterations in proliferation or cell death were noted. On the other hand, analysis of global gene expression changes in *Cables2*-deficient EpiLC showed enrichment of genes associated with “p53 signalling pathway” and “negative regulation of cell proliferation” among upregulated genes in *Cables2*-deficient EpiLCs and genes associated with “pluripotent of stem cells” and “positive regulation of cell proliferation” among downregulated genes in *Cables2*-deficient EpiLCs. Moreover, the marked downregulation of *Nanog* expression in EpiLCs was confirmed *in vivo* and reporter assays in EpiLCs indicated *Nanog* expression is *Cables2*-dependent. Given that epiblast expression of *Nanog* is known to be Nodal/Smad2-dependent (Sakaki-Yumoto et al., 2013; Sun et al., 2014), it is possible that *Cables2* in the epiblast interacts with this pathway to ensure *Nanog* expression. The possibility that loss of *Cables2* causes some dysregulation of pluripotency and/or proliferation-related genes, including *Nanog* and *Cdkn1a/p21*, in epiblast after E5.5 warrants further investigation.

To data, there is no report addressing the molecular function of Cables2 *in vitro* and *in vivo*. Analysis of the Cables2 protein sequence using publicly available protein domain prediction tools fails to predict any enzymatic, DNA-binding, nor transcription regulatory domains. Thus, we speculate that Cables2 functions as a bridging factor, or scaffold protein, mediating interaction between transcription regulatory factors. A similar role for Cables1 in connecting CDKs and nonreceptor tyrosine kinases has been proposed. Indeed, our co-IP experiments showed the interaction of Cables2 with transcription regulatory factors, β-catenin, Smad2 and Oct4, which are known to form a protein complex to activate the promoter of common target genes (**Figure 4B, 5L and Figure 6—figure supplement 2**) (Funa et al., 2015). We also demonstrated that Cables2 interacted with Smad2 in nuclear and cytoplasm (**Figure 5M and N**). These results suggest that Cables2 functions as a nuclear cofactor for Smad2 although the mechanism underlying enhancement of Smad2 transcriptional activity by Cables2 remains to be elucidated. Interestingly, extracellular signal-regulated kinase (ERK) promotes Smad2 transcriptional activity, but suppresses Smad3 transcriptional activity by phosphorylation of their linker region in helper T cells (Chang et al., 2011; Funaba et al., 2002; Yoon et al., 2015b). The linking function of Cables2 could promote complex formation between Smad2 and other transcription factors to form a transcription complex or it could provide a platform for posttranslational modifiers such as ERK.

In conclusion, *Cables2* plays an essential role in mouse embryogenesis and targeted disruption of *Cables2* leads to post-gastrulation embryonic lethality. *Cables2* is required in both the VE to promote axis formation and in the epiblast to promote continued growth of the embryo. Our investigations support a role for Cables2 in one or more of the signalling pathways crucial for the development of the peri-implantation stage mouse embryo since it can physically interact with both β-catenin and Smad2. Cables2 may serve as a scaffold protein that facilitates a variety of molecular interactions. Given the pleiotropic nature of the *Cables2* null phenotype it is likely that further genetic and molecular analyses will uncover additional roles for the protein.

## METHODS

### Animals and husbandry

ICR mice were purchased from CLEA Japan Co. Ltd. (Tokyo, Japan); C57BL/6N mice were purchased from Charles River Laboratory Japan Co. Ltd (Yokohama, Japan). For production of staged embryos, the day of fertilization as defined by the appearance of a vaginal plug was considered to be embryonic day 0.5 (E0.5). Animals were kept in plastic cages (4 – 5 mice per cage) under specific pathogen-free conditions in a room maintained at 23.5°C ± 2.5°C and 52.5% ± 12.5% relative humidity under a 14-h light:10-h dark cycle. Mice had free access to commercial chow (MF; Oriental Yeast Co. Ltd., Tokyo, Japan) and filtered water throughout the study. Animal experiments were carried out in a humane manner with approval from the Institutional Animal Experiment Committee of the University of Tsukuba in accordance with the Regulations for Animal Experiments of the University of Tsukuba and Fundamental Guidelines for Proper Conduct of Animal Experiment and Related Activities in Academic Research Institutions under the jurisdiction of the Ministry of Education, Culture, Sports, Science, and Technology of Japan.

### Generation of *Cables2*-deficient mice

The targeted ES cell clone *Cables2*^tm1(KOMP)Vlcg^ was purchased from KOMP (project ID: VG1608, clone number: 16085A-D3). To generate *Cables2*-deficient mice, ES cells were aggregated with the wild-type morula and transferred to pseudopregnant female mice. Male chimeras that transmitted the mutant allele to the germ line were mated with wild-type females to produce *Cables2*-deficient mice with the C57BL/6N background. Adult mice were genotyped using genomic DNA extracted from the tail. For whole-mount *in situ* hybridization, embryos were genotyped using a fragment of yolk sac and Reichert membrane. Samples were dispensed into lysis solution (50 mM Tris-HCl, pH 8.5, 1 mM EDTA, 0.5% Tween 20) and digested with proteinase K (1 mg/mL) at 55°C for 2 hours, inactivated at 95°C for 5 minutes, and then subjected to PCR. For paraffin slides, embryos were genotyped using tissue picked from sections and digested directly with proteinase K (2 mg/mL) in PBS. For others experiments, after collecting data, the whole embryos were used for genotyping. Genotyping PCR was performed with AmpliTag Gold 360 Master Mix (Thermo Fisher Scientific K.K., Tokyo, Japan) using the following primers: *Cables2* D3-1: 5’-ACTGCAGAAGCTGGAGGAAA-3’; *Cables2* D3-2: 5’-TCAAGGTGTCTGCCCTATCC-3’; *Cables2* D3-3: 5’-AGGGGATCCGCTGTAAGTCT-3’.

### *Nanog*-GFP reporter mice

*Nanog*-GFP transgenic mice (RBRC02290) were obtained from Riken BioResource Center (BRC; Tsukuba, Japan). Animals were kept and maintained under the same conditions as described above. To produce the *Nanog*-GFP reporter in the homozygous *Cables2* background, *Cables2* heterozygotes were first crossed with *Nanog^GFP/+^* to obtain *Cables2^+/-^:Nanog^GFP/+^*. Subsequently, to obtain *Cables2^-/-^:Nanog^GFP/+^* embryos, *Cables2* heterozygotes were mated with *Cables2^+/-^:Nanog^GFP/+^* mice and the embryos were collected at E6.5 or E5.5. All embryos were then genotyped using both *Cables2* genotyping primers and the GFP following primers: GFP F: 5’-ACGTAAACGGCCACAAGTTC-3’; GFP R: 5’-TGCTCAGGTAGTGGTTGTCG-3’.

### TOPGAL reporter mice

B6.Cg-Tg(TOPGAL) transgenic mice carrying LEF/TCF reporter of Wnt/β-catenin signalling were used for visualizing Wnt signalling pathway *in vivo*. TOPGAL mice were obtained from Riken BRC (RBRC02228). Animals were kept and maintained under the same conditions as described above. To produce the TOPGAL reporter in the homozygous *Cables2* background, TOPGAL heterozygotes were crossed with *Cables2* heterozygotes subsequently and finally, homozygous *Cables2* carrying TOPGAL transgene were collected at E7.5 together with littermates. All embryos were stained S-gal (Sundararajan et al., 2012) and then genotyped using both *Cables2* genotyping primers and TOPGAL following primers: TOPGAL-TK F: 5’-CGAGGTCCACTTCGCATATT-3’; LacZ R: 5’-TATTGGCTTCATCCACCACA-3’.

### Cell culture

NIH3T3, Cos-7, 293T cells were obtained from The American Type Culture Collection (Manassas, Virginia) and 293FT cells were purchased from Thermo Fisher Scientific. These cells were authenticated by the suppliers and no mycoplasma contamination was detected by DAPI staining. Cells were cultured in Dulbecco’s modified Eagle’s medium (DMEM) supplemented with 10% heat-inactivated fetal bovine serum. Mouse embryonic stem cells (ESCs) were maintained on 0.1% gelatine-coated dishes in mouse ESC medium consisting of DMEM containing 20% knockout serum replacement (KSR; Thermo Fisher Scientific), 1% non-essential amino acids (Thermo Fisher Scientific), 1% GlutaMAX (Thermo Fisher Scientific), 0.1 mM 2-mercaptoethanol (Thermo Fisher Scientific), and leukemia inhibitory factor (LIF)-containing conditioned medium, supplemented with two chemical inhibitors (2i), i.e., 3 μM CHIR99021 (Stemgent inc., Cambridge, Massachusetts) and 1 μM PD0325901 (Stemgent). The epiblast-like cells (EpiLCs) were induced by plating 2.0 × 10^5^ ESCs on human fibronectin (Corning inc., Corning, New York)-coated 6-well plates in N2B27-containing NDiff 227 medium (Takara Bio Inc., Shiga, Japan) supplemented with 20 ng/mL activin A, 12 ng/mL bFGF, and 1% KSR (Guo et al., 2009). All cells were cultured in an atmosphere of 5% CO_2_ at 37°C.

### RT-PCR, RT-qPCR and RNA-seq

Cultured ES cells, about 130 blastocysts, and 21 embryos at E7.5 were collected. Total RNAs from blastocysts and embryos were extracted using Isogen (Nippon Gene Co., Ltd., Tokyo, Japan). RNA from ESCs was collected using an RNeasy Mini Kit (Qiagen K.K., Tokyo, Japan). The cDNA was synthesized using Oligo-dT primer (Thermo Fisher Scientific) and SuperScript III Reverse Transcriptase (Thermo Fisher Scientific) in a 20-μL reaction mixture. The primers were: *Cables2* F: 5’-CACCAGCTGGCACAGAACTA-3’; *Cables2* R: 5’-GCTTGAGGATCAAGTGTGGTTCAAAGTC-3’; Glyceraldehyde-3-phosphate dehydrogenase (*Gapdh*) F: 5’-ACCACAGTCGATGCCATCAC-3’; *Gapdh* R: 5’-TCCACCACCCTGTTGCTGTA-3’.

RT-qPCR was performed using SYBR Premix Ex Taq II (Takara) and the Thermal Cycler Dice Real Time System (Takara) according to the manufacturer’s instructions. The *Nanog* gene expression level was normalized to the endogenous *Gapdh* expression level. The primers used were: *Nanog* F: 5’-CCTGAGCTATAAGCAGGTTAAG-3’; *Nanog* R: 5’-GTGCTGAGCCCTTCTGAATC-3’; *Gapdh* qPCR F: 5’-TGGAGAAACCTGCCAAGTATG-3’; *Gapdh* qPCR R: 5’-GGAGACAACCTGGTCCTCAG-3’.

RNA sequencing analysis was performed by Tsukuba i-Laboratory LLP as previously described (Ohkuro et al., 2018). Briefly, total RNAs were extracted from wild-type and *Cables2*-deficient EpiLCs at 2 days post-induction (*n* = 3) using RNeasy Plus Mini Kit (Qiagen). RNA quality was evaluated using Agilent Bioanalyzer with RNA 6000 Pico kit (Agilent Technologies Japan, Ltd., Tokyo, Japan). An amount of 500 ng total RNA was used for RNA-seq library preparation with NEB NEBNext rRNA Depletion Kit and ENBNext Ultra Directional RNA Library Prep Kit (New England Biolabs Japan Inc., Tokyo, Japan); 2 × 36 base paired-end sequencing was performed with NextSeq500 (Illumina K.K., Tokyo, Japan) by Tsukuba i-Laboratory LLP (Tsukuba, Japan). The RNA-seq data have been deposited in the NCBI GEO database (accession no. GSE120366). The DAVID Bioinformatics Resources was used for GO terms and KEGG pathway enrichment analyses of differentially expressed genes with at least 2-fold change and FDR < 0.05.

### Vector construction

Part of *Cables2* cDNA containing exons 1 and 2 was cloned in-frame into pBlueScript KS+ at the *Bam*HI site, and the fragment containing exons 3 – 10 was cloned into the pcDNA3 vector at the *Bam*HI site. These fragments were obtained and amplified from a mouse embryo E7.5 cDNA library and sequenced. The part covering *Cables2* exons 1 and 2 was cut at the *Afe*I site and ligated into the pcDNA3 vector containing exons 3 – 10. The full-length *Cables2*, FLAG-tagged *Cables1*, FLAG-tagged *Cables2*, and mouse Kpna (Importin α) NLS-fused FLAG-tagged *Cables2* genes were cloned into the *Eco*RI site of pCAG vector. The full-length *Smad2* and *Pou5f1* cDNA was amplified by PCR from mouse B6 ESC cDNA library and cloned into the pCAG vector. A 1.5-kb *Cables2* riboprobe was prepared by amplification from the full-length cDNA template with the pcDNA3 backbone, synthesized with Sp6 polymerase, and labelled with digoxigenin as a riboprobe.

A ROSA26 knock-in vector was constructed by insertion of CAG-Venus-IRES Pac gene expression cassette (Khoa et al., 2016) into the entry site of pROSA26-1 vector (kindly gifted from Philippe Soriano, Addgene plasmid # 21714) (Soriano, 1999). The *Cables2^-/-^; ROSA^YFP/+^* was generated by electroporation of the ROSA26 knock-in vector (pROSA26-CAG-Venus-IRES Pac) into *Cables2^-/-^* ESCs. The CAG-tdTomato-2A and 3xFLAG sequences were inserted in the upstream and downstream of *Cables2* cDNA, respectively, to make CAG-tdTomato-2A-Cables2-3xFLAG vector. The expression of tdTomato and FLAG-tagged Cables2 in the CAG-tdTomato-2A-Cables2 vector-transfected 293FT cells were evaluated by fluorescent microscopy and western blot analysis with anti-FLAG antibody, respectively (data not shown).

### Production of *Cables2* rescue chimeras by Tetraploid complementation assay

Tetraploid (4n) wild-type embryos were made by electrofusing diploid (2n) embryos at two-cell-stage and cultured up to morula stage. The 4n wild-type morula were aggregated with *Cables2^-/-^; ROSA^YFP/+^* or *Cables2^-/-^; ROSA^YFP/+^; CAG-tdTomato-2A-Cables2-3xFLAG* ESCs to form blastocyst chimeras. B6N wild-type ESC was used as a control for tetraploid complementation assay. To obtain comparable control embryos at each stage of development, an equal number of control blastocyst chimeras were transferred together with *Cables2^-/-^* blastocyst chimeras to a pseudopregnant recipient mouse at E2.5. Embryos were recovered at from E6.5 to E9.5 and the contribution of ESCs was evaluated by YFP or tdTomato fluorescence signals.

### Whole-mount *in situ* hybridization (WISH)

All embryos were dissected from the decidua in PBS with 10% fetal bovine serum and staged using morphological criteria (Downs and Davies, 1993) or described as the number of days of development. WISH was carried out following standard procedures, as described previously (Rosen and Beddington, 1994). Briefly, embryos were fixed overnight at 4°C in 4% paraformaldehyde in PBS, dehydrated, and rehydrated through a graded series of 25% – 50% – 75% methanol/PBS. After proteinase K (10 μg/mL) treatment for 15 minutes, embryos were fixed again in 0.1% glutaraldehyde/4% paraformaldehyde in PBS. Pre-hybridization at 70°C for at least 1 hour was conducted before hybridization with 1–2 μg/mL digoxigenin-labelled riboprobes at 70°C overnight. Pre-hybridization solution included 50% formamide, 4×SSC, 1% Tween-20, heparin (50 μg/mL) (Sigma-Aldrich Japan K.K, Tokyo, Japan) and hybridization was added more yeast RNA (100 μg/mL) and Salmon Sperm DNA (100 μg/mL) (Thermo Fisher Scientific). For post-hybridization, embryos were washed with hot solutions at 70°C including 50% formamide, 4×SSC, 1% SDS, and treated with 100 μg/mL RNase A at 37°C for 1 hour. After additional stringent hot washes at 65°C including 50% formamide, 4×SSC, samples were washed with TBST, pre-absorbed with embryo powder, and blocked in blocking solution (10% sheep serum in TBST) for 2–5 hours at room temperature. The embryo samples were subsequently incubated with anti-digoxigenin antibody conjugated with alkaline phosphatase anti-digoxigenin-AP, Fab fragments (Roche Diagnostics K.K., Tokyo, Japan) overnight at 4°C. Extensive washing in TBST was followed by washing in NTMT and incubation in NBT/BCIP (Roche) at room temperature (RT) until colour development. After completion of in situ hybridization (ISH), embryos were de-stained in PBST for 24 – 48 hours and post-fixed in 4% paraformaldehyde in PBS. Embryos were processed for photography through a 50%, 80%, and 100% glycerol series. Before embedding for cryosectioning, embryos were returned to PBS and again post-fixed in 4% paraformaldehyde in PBS. The specimens were placed into OCT cryoembedding solution, flash-frozen in liquid nitrogen, and cut into sections 14 μm thick using a cryostat (HM525 NX; Thermo Fisher Scientific). The following probes were used for WISH: *Bmp4* (Jones et al., 1991), *Brachyury (T)* (Herrmann, 1991), *Cer1* (Belo et al., 1997), *Foxa2* (Sasaki and Hogan, 1993), *Fgf8* (Bachler and Neubüser, 2001), *Lefty1/2* (Meno et al., 1996), *Lhx1* (Shawlot and Behringer, 1995), *Nanog* (Chambers et al., 2003), *Nodal* (Conlon et al., 1994), *Oct4* (Schöler et al., 1990), *Otx2* (Simeone et al., 1993), *Sox2* (Avilion et al., 2003), *Sox17* (Kanai et al., 1996), and *Wnt3* (Roelink et al., 1990).

### Co-immunoprecipitation (Co-IP)

At 1 day before transfection, aliquots of 5 × 10^4^ 293FT cells were seeded onto poly-L-lysine (PLL)-coated 6-cm dishes and co-transfected with 2 μg of each pCAG-based expression vector using Lipofectamine 3000 (Thermo Fisher Scientific). After 48 hours, the cells were washed once with PBS, resuspended in RIPA buffer (50 mM Tris-HCl, pH 7.4, 150 mM NaCl, 1 mM EDTA, 1% deoxycholic acid and 1% Nonidet P-40 [NP-40]) containing protease inhibitor cocktail (Roche Diagnostics) and placed on ice for 30 minutes. The supernatant was collected after centrifugation and incubated with Dynabeads Protein G (Veritas Co., Tokyo, Japan) and mouse anti-DYKDDDDK (FLAG)-tag antibody (KO602-S; TransGenic Inc., Fukuoka, Japan) overnight at 4°C. The beads were washed four times with PBS, resuspended in Laemmli sample buffer, and boiled. The precipitated proteins were analysed by SDS-polyacrylamide gel electrophoresis (SDS-PAGE) and western blotting using the ECL Select Western Blotting Detection System (GE Healthcare Japan Co., Ltd., Tokyo, Japan) and a LAS-3000 imaging system (GE Healthcare). The FLAG antibody was then washed out and the membrane was re-stained with anti-β-catenin antibody (#8480, Cell Signalling Technology), anti-HA antibody (3F10, Roche), anti-Smad2 antibody (#5339, Cell Signalling Technology) or anti-GAPDH antibody (sc-25778, Santa Cruz).

### Cell fractionation

One day after seeding of 3 × 10^6^ 293FT cells on 10 cm dishes, the cells were co-transfected with 2.5 μg each of HA-Smad2 and FLAG-Cables2 expression vectors using Lipofectamine 3000 (Thermo Fisher Scientific). After 24 hours, the cells were washed once with PBS, scraped from dishes and then collected using NE-PER Nuclear and Cytoplasmic Extraction Reagents (Thermo Fisher Scientific). One-third of the cell suspension in 0.1% NP-40 was used as whole cell lysate. The whole cell suspension and nuclear fraction were sonicated and added the equal volume of RIPA buffer. Aliquots of each protein lysate were applied for co-IP with Dynabeads Protein G and mouse anti-FLAG M2 antibody (F1804) or rat anti-HA antibody (3F10) and the fractionation were confirmed by expression of α Tubulin (sc-5286, Santa Cruz) and PARP-1 (sc-8007, Santa Cruz).

### Luciferase reporter assay

A total of 50,000 cells were plated in PLL-coated 96-well tissue culture plates. After overnight culture, the cells were transfected with a specific promoter-driven firefly reporter plasmid and *Renilla* luciferase control plasmid, pRL-TK, using Lipofectamine 3000 (Thermo Fisher Scientific) and opti-MEM (Thermo Fisher Scientific). Luciferase activity was analysed using a luminometer and a Dual-Glo Luciferase assay kit according to the manufacturer’s instructions (Promega K.K., Tokyo, Japan). The firefly luciferase values were normalized to those of *Renilla* luciferase. To evaluate β-catenin activity, cells were transiently transfected with TOPflash (TOP) or FOPflash (FOP) reporter plasmids carrying multiple copies of a wild-type or mutated TCF-binding site, respectively. Relative activity was calculated as normalized relative light units of TOPflash divided by normalized relative light units of FOPflash. To examine the SMAD2 activity, cells were transfected with ARE-luc reporter plasmid, which expresses firefly luciferase driven by a SMAD2- and FAST1-dependent promoter, pRL-TK, a FAST1 expression plasmid (Hayashi et al., 1997), and a constitutive-active mutant form of ALK5 expression plasmid. The (CAGA)_9_-luc reporter plasmid was used with pRL-TK and a constitutive-active mutant form of ALK5 expression plasmid to evaluate the effect of Cables1 or 2 on the SMAD3 activity in 293FT cells (Dennler et al., 1998). *Nanog* promoter activity was evaluated using the Nanog5p-luc reporter plasmid, which contains 2.5 kb of 5’ promoter region of the mouse *Nanog* gene. Nanog5p reporter was a gift from Austin Cooney (Addgene plasmid #16337). The experiment was performed in triplicate and repeated at least three times. Statistical analysis was performed using the Mann-Whitney U-test. Two-tailed *P*-values at less than 0.05 were considered as statistically significant.

### Indirect immunofluorescence assay (IFA)

After 24 or 48 hours transfection, cells were washed twice with PBS and then fixed with 4% paraformaldehyde in PBS for 10 minutes. Permeabilization of cell membranes were done with 0.1% Triton X-100 in PBS for 20 minutes or methanol for 5 minutes. After blocking with 10% goat serum or Superblock blocking buffer (Thermo Fisher Scientific) for 30 minutes, cells were incubated overnight at 4°C with mouse anti-FLAG antibody. Then, the cells were washed with PBS and incubated with Alexa Fluor-conjugated anti-mouse IgG antibody. Fluoresence signals were detected using a BZ-X700 fluorescent microscope (Keyence Co., Ltd., Osaka, Japan).

### Histology, EdU, and TUNEL assay

Mouse uteri including the decidua were collected and fixed in 4% paraformaldehyde in PBS. Subsequently, paraffin blocks were made by dehydration in ethanol, clearing in xylene, and embedding in paraffin. Embryo sections 5 μm thick were cut (Microm HM 335E; Thermo Fisher Scientific) and placed on glass slides (Matsunami Glass Ind., Ltd., Osaka, Japan). For haematoxylin-eosin (HE) staining, slides were deparaffinized and rehydrated through an ethanol series, and then stained with HE.

To label the proliferating embryonic cells, pregnant mice at E6.5 were injected intraperitoneally with 5-ethynyl-2’-deoxyuridine (EdU) at 200 μL/mouse and sacrificed 4 hours later. Embryos were embedded in paraffin blocks, and sections were refixed in 4% paraformaldehyde and permeabilized in 0.5% Triton X-100/PBS. EdU assay was performed with a Click-iT Plus EdU Imaging Kit (Thermo Fisher Scientific) and TUNEL assay was performed with a Click-iT Plus TUNEL Assay for In situ Apoptosis Detection kit (Thermo Fisher Scientific) according to the manufacturer’s protocol. As the final step, embryo sections were co-stained with Hoechst 33342, observed under a microscope (BZ-X700; Keyence), and cell number was counted using ImageJ software.

## Acknowledgements

We thank all members of the Sugiyama Laboratory and Laboratory Animal Resource Center for helpful discussions and encouragement. Furthermore, we are indebted to T. Chiba, K. Kako, H. Katayama, and Y. Yuda for discussion and comments on this manuscript.

## Figure supplements

**Figure 1—figure supplement 1.**
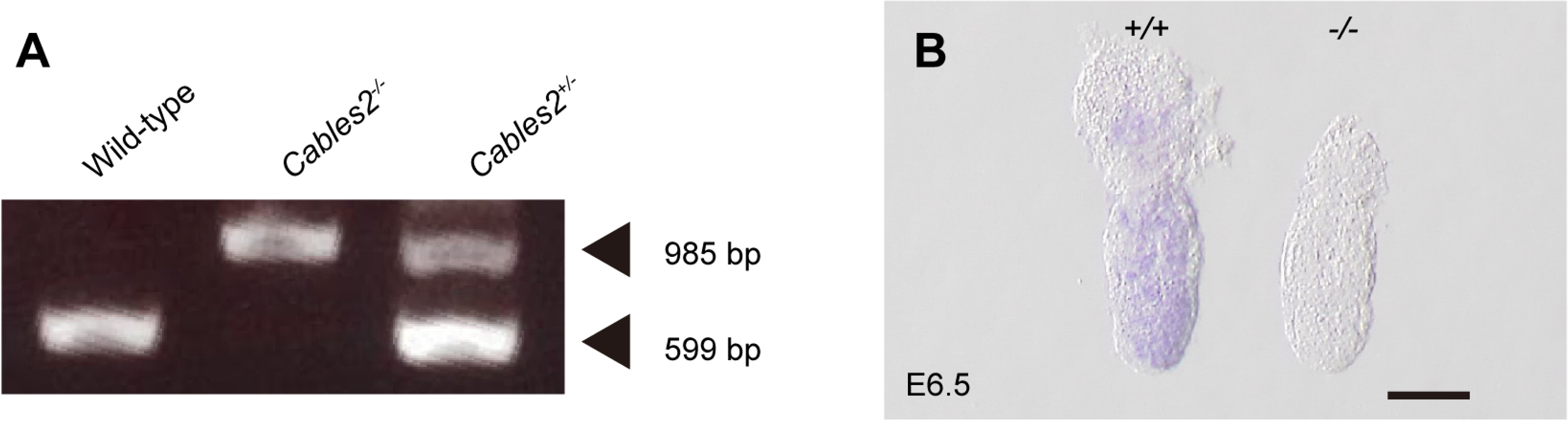
Genotyping and expression of *Cables2*. (A) Genotyping by PCR analysis from 5 whole embryo samples using three primers. Bands at 985 bp and 599 bp represent mutant and 6 wild-type *Cables2* alleles, respectively. (B) At E6.5, *Cables2* was expressed ubiquitously in 7 both extra- and embryonic parts in wild-type (left), in comparison with homozygous mutants 8 (right). Antisense *Cables2* probe was used for WISH to confirm that *Cables2*-deficient 9 embryos lacked expression. Scale bar, 100 μm.

**Figure 2—figure supplement 1.**
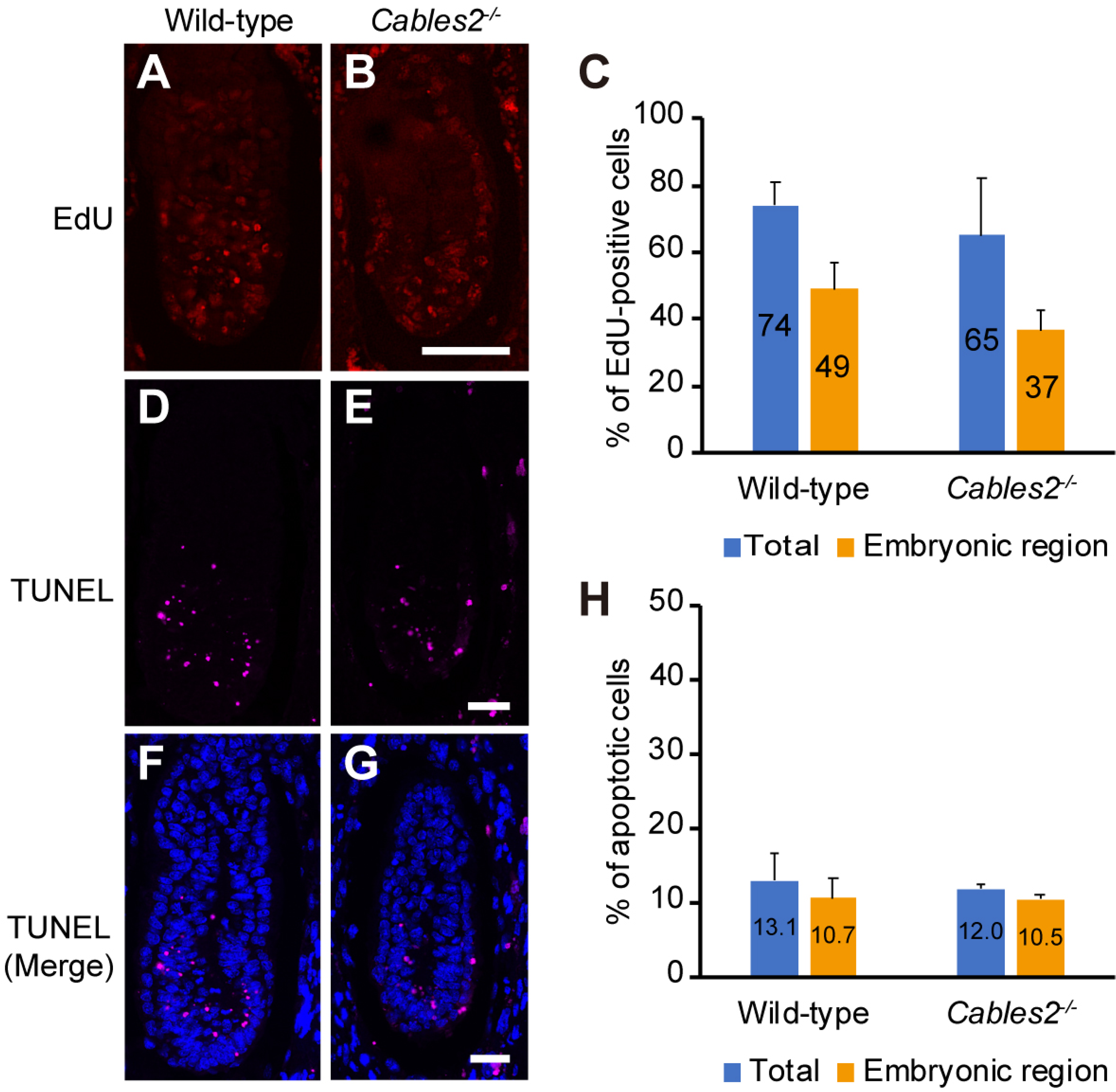
Proliferating and apoptotic cells in E6.5 *Cables2* mutant embryos. (A–C) The EdU-incorporating cells represented the proliferation of cells in both wild-type and Cables-deficient embryos (*n* = 4). The proliferated cells were counted in whole embryo (Total) or only in embryonic region. (D–F) Apoptotic cells were detected in both wild-type and *Cables2*-deficient embryos (*n* = 3), in whole embryo or in embryonic part. The average percentage was calculated by number of counted cells normalized to total number of cells within the embryo. At least 2 slides were counted per embryo. Y error bars: Standard of deviation (SD). Statistical significance was determined using Student’s t test (*P* < 0.05). Scale bars, 50 μm.

**Figure 4—figure supplement 1.**
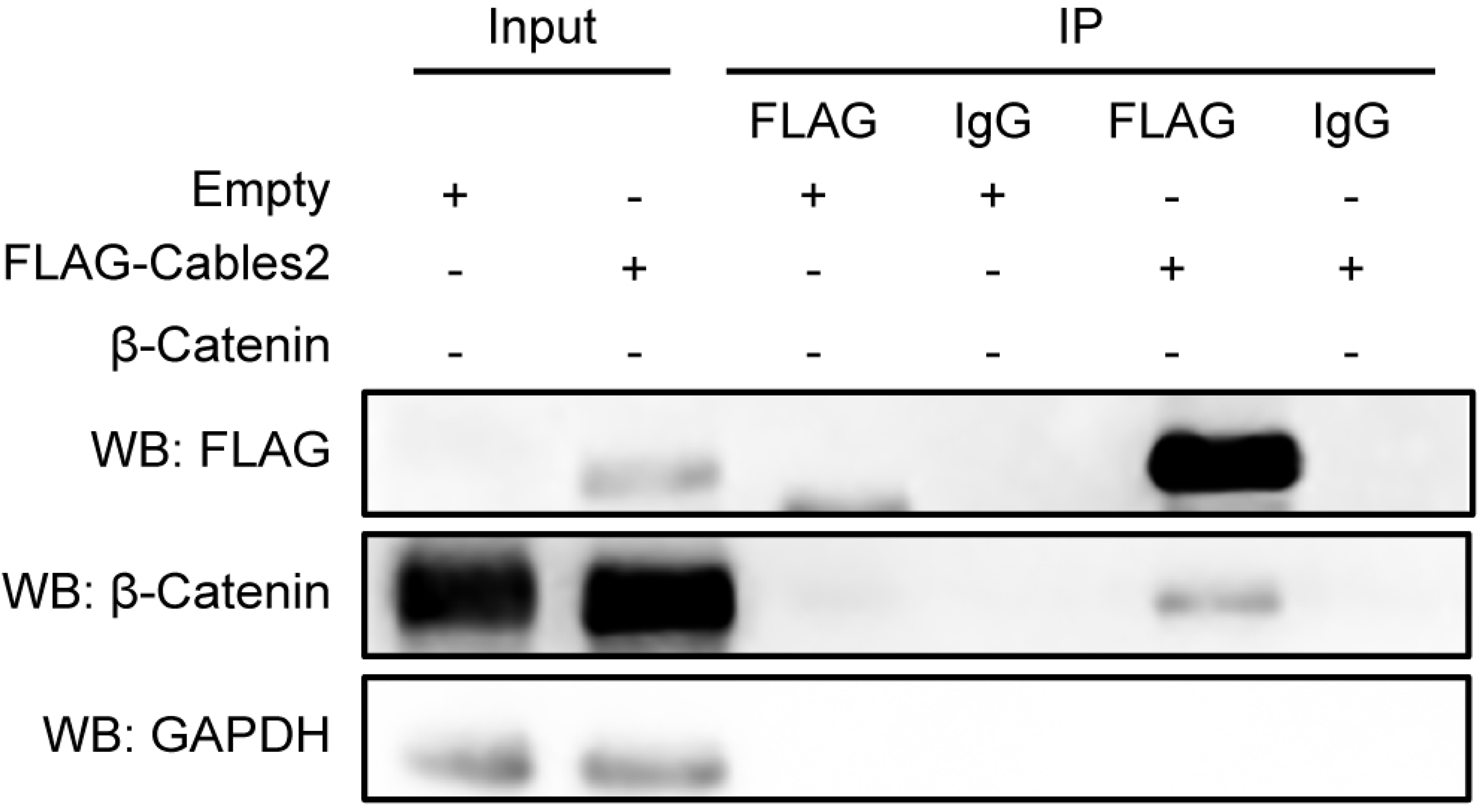
Interaction of Cables2 with endogenous β-catenin. Co-IP showing the physical interaction of FLAG-Cables2 with endogenous β-catenin in 293FT cells. Anti-GAPDH antibody was used as a negative control for evaluating specific interaction. Experiment was repeated at least twice and reliably reproduced.

**Figure 5—figure supplement 1.**
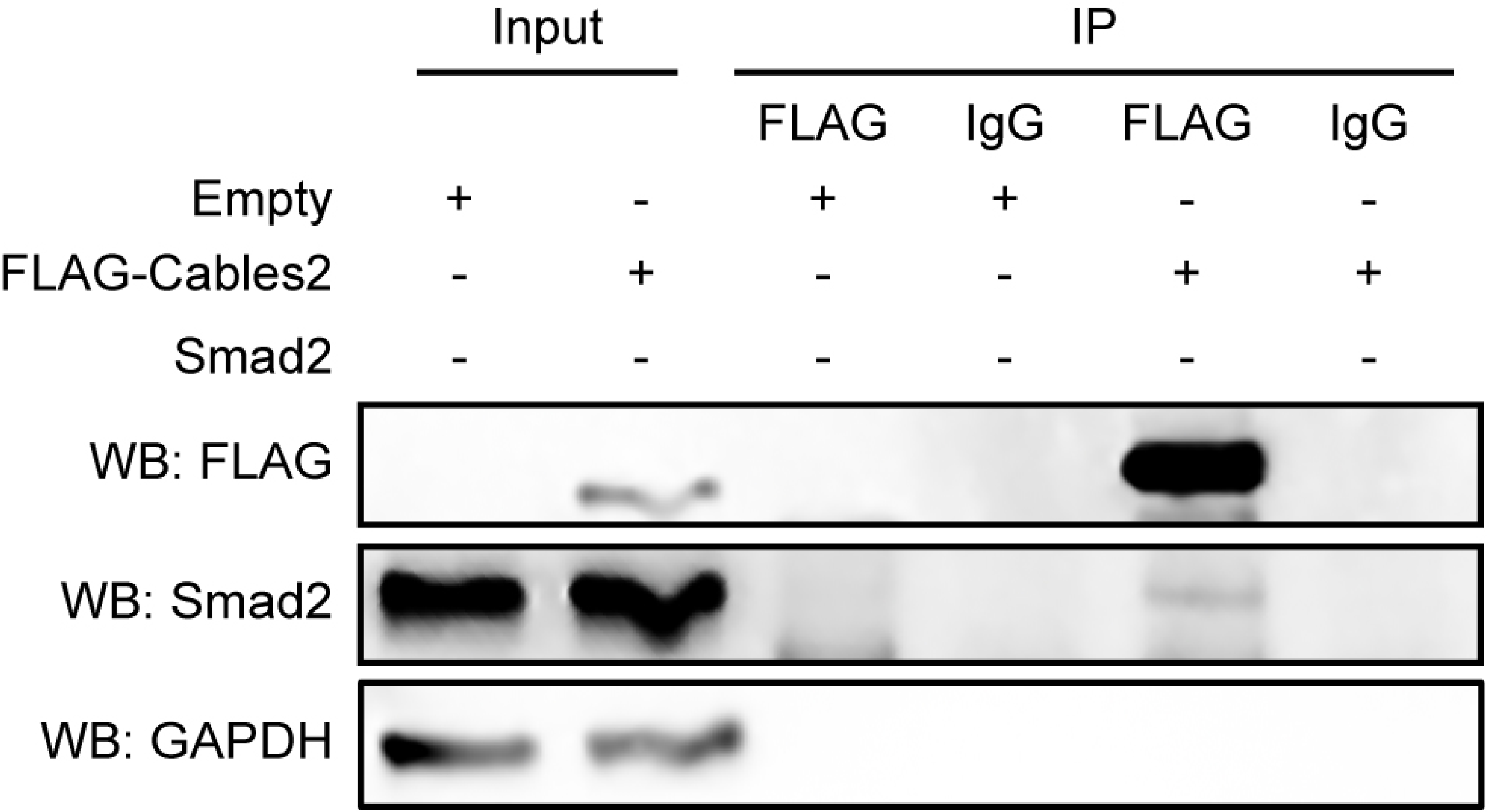
Interaction of Cables2 with endogenous Smad2. Co-IP showing the physical interaction of FLAG-Cables2 with endogenous Smad2 in 293FT cells. Anti-GAPDH antibody was used as a negative control for evaluating specific interaction. Experiment was repeated at least twice and reliably reproduced.

**Figure 6—figure supplement 1.**
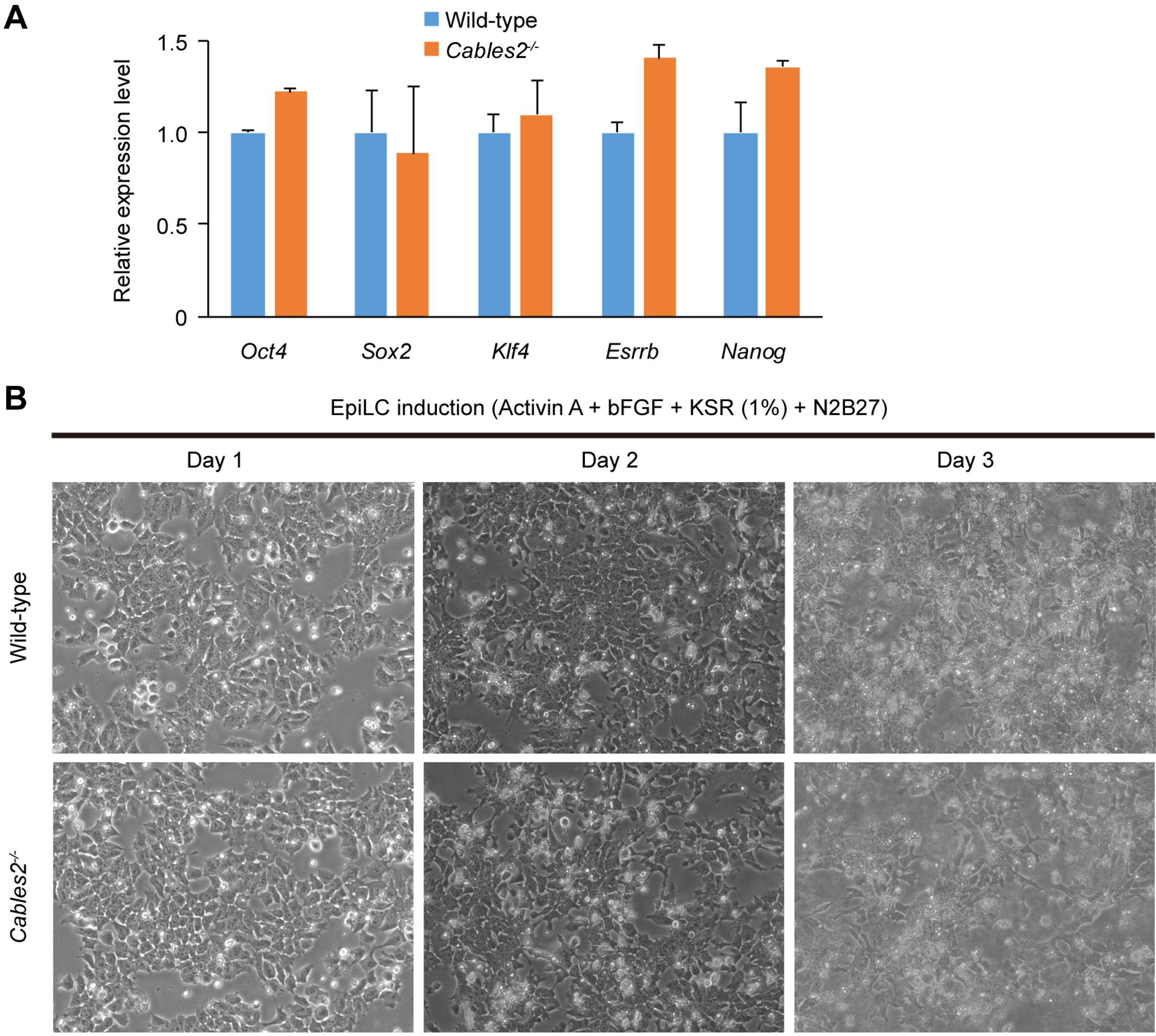
EpiLC induction from *Cables2^-/-^* ESCs and expression of pluripotency genes. (A) RT-qPCR showing the expression levels of pluripotency genes in wild-type and *Cables2^-/-^* ESCs. (B) Wild-type and *Cables2^-/-^* ESCs were maintained with 2i/LIF and induced to differentiate into EpiLCs within 3 days. No differences were observed between wild-type and *Cables2^-/-^* EpiLCs in cell proliferation or cell morphology.

**Figure 6—figure supplement 2.**
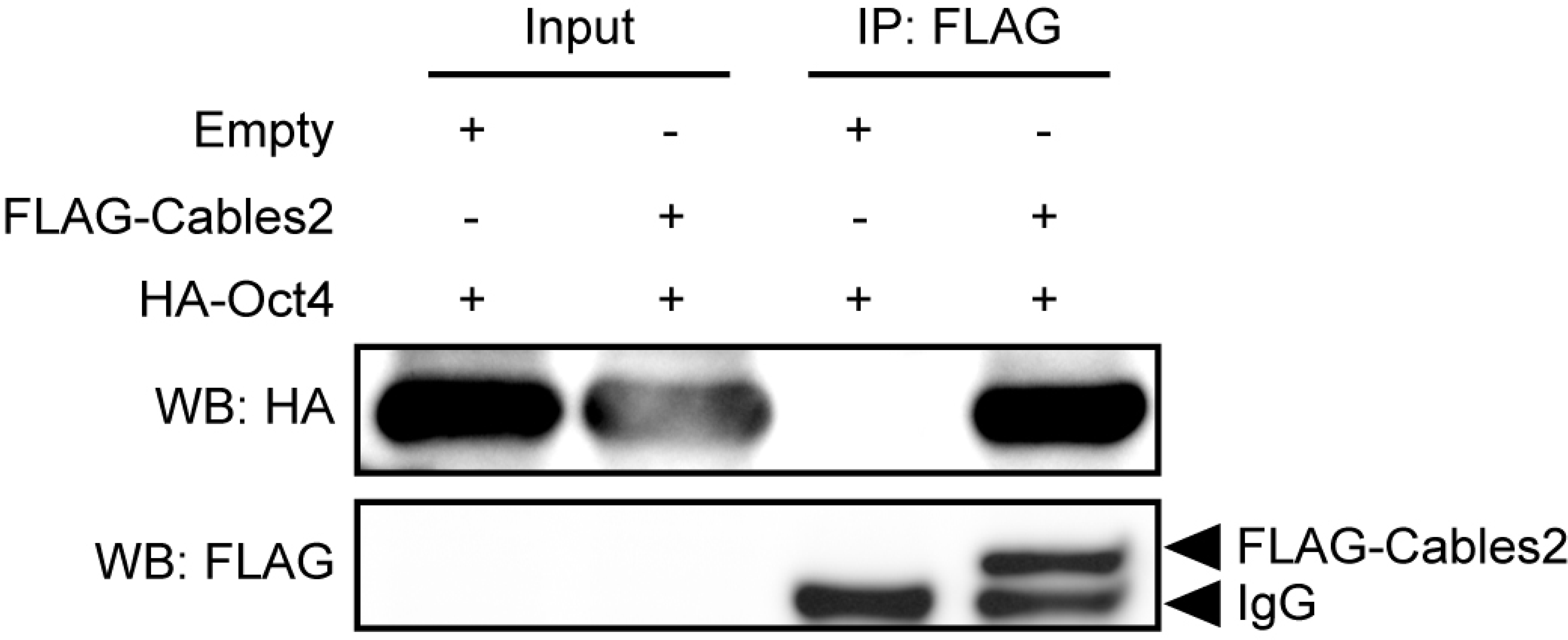
Interaction of Cables2 with exogenous HA-Oct4. Co-IP showing the physical interaction of FLAG-Cables2 with HA-Oct4 in 293FT cells. Anti-GAPDH antibody was used as a negative control for evaluating specific interaction. Experiment was repeated at least twice and reliably reproduced.

**Figure 6—figure supplement 3.**
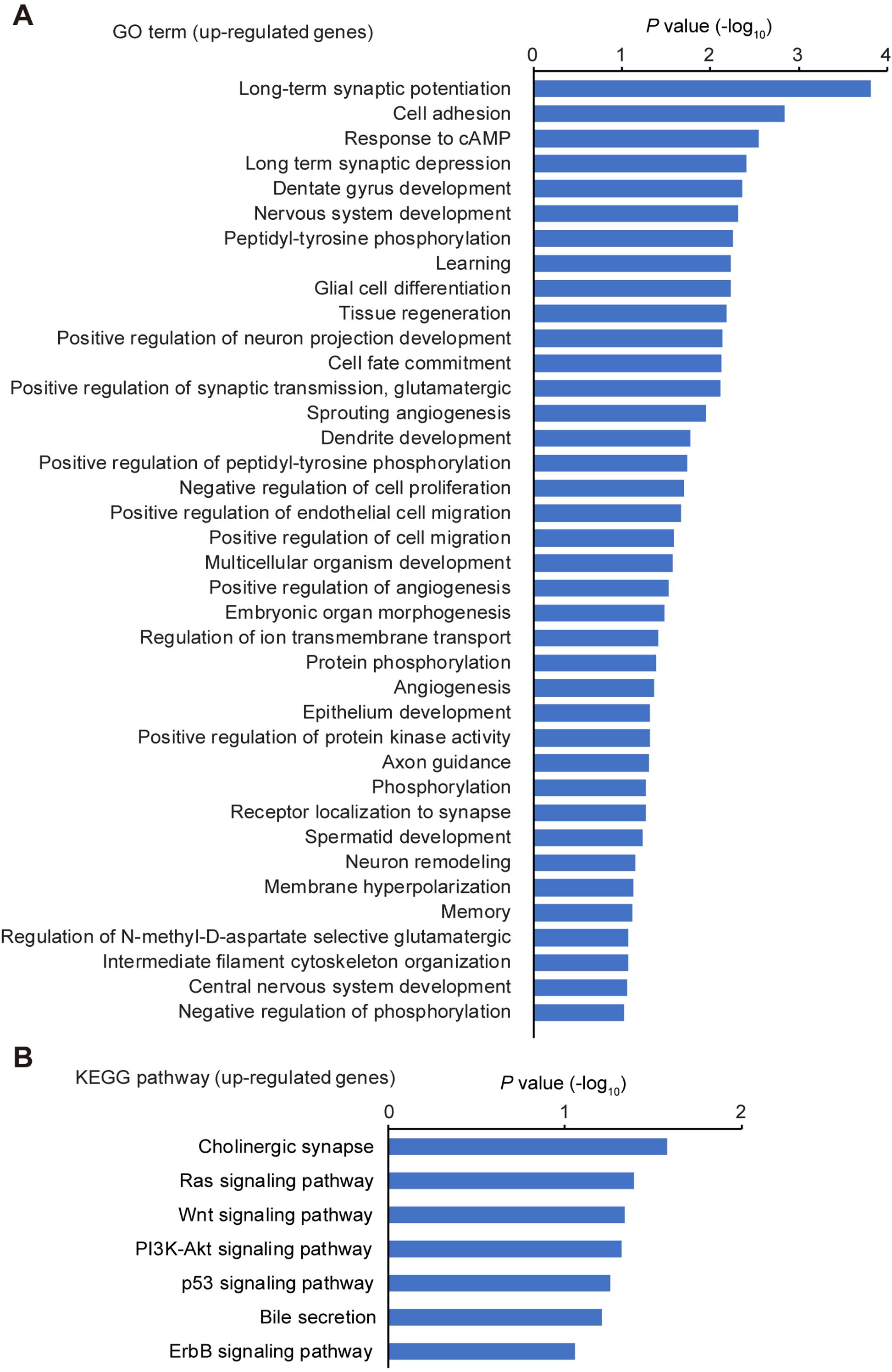
Gene enrichment analyses of up-regulated genes by the loss of Cables2. (A, B) GO term (A) and KEGG pathway (B) enrichment analyses of 122 up-regulated genes in *Cables2*-deficient EpiLCs compared with wild-type EpiLCs.

**Figure 6—figure supplement 4.**
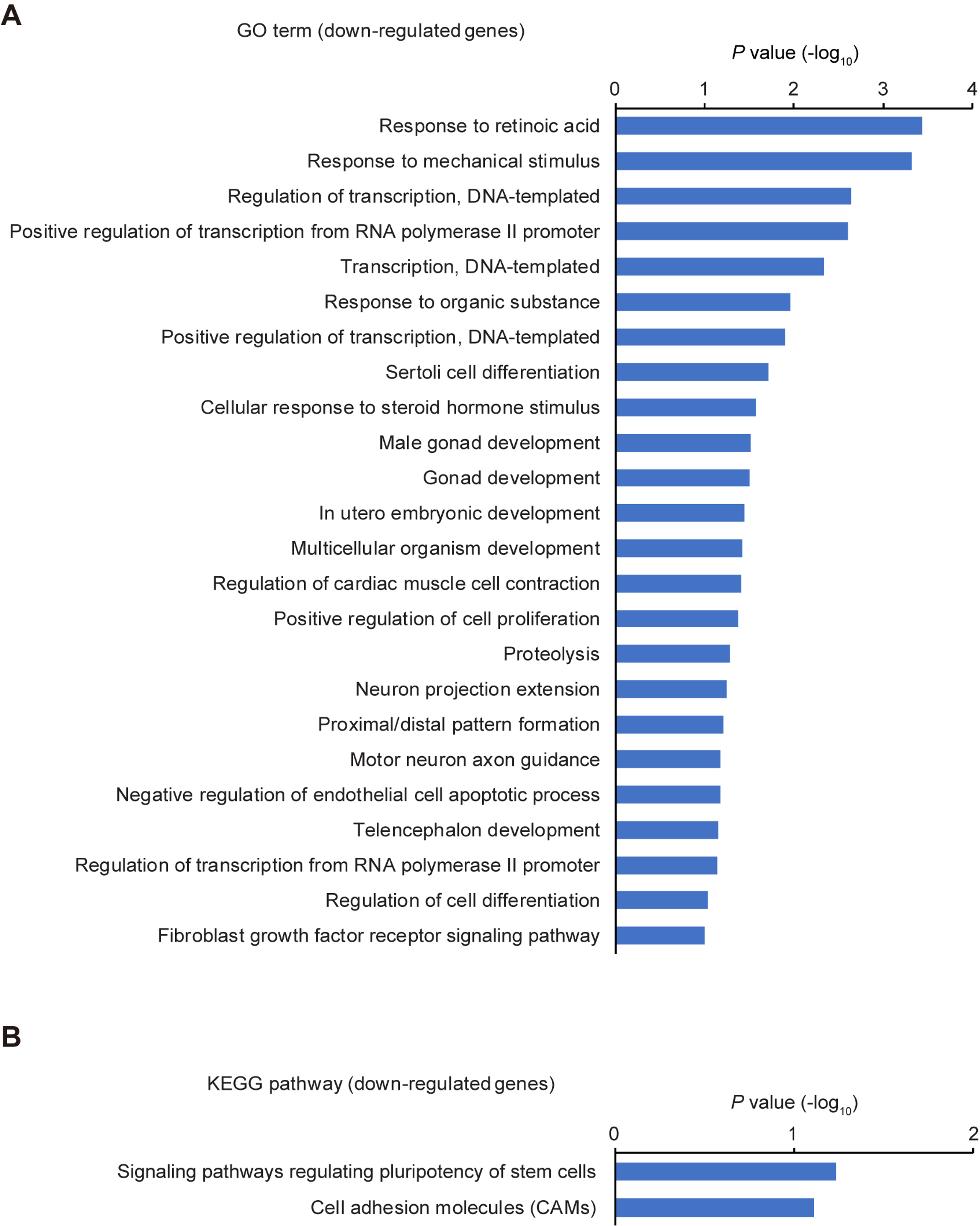
Gene enrichment analyses of down-regulated genes by the loss of Cables2. (A, B) GO term (A) and KEGG pathway (B) enrichment analyses of 52 down-regulated genes in *Cables2*-deficient EpiLCs compared with wild-type EpiLCs.

